# Thiamine pyrophosphokinase deficiency induces Alzheimer’s pathology

**DOI:** 10.1101/2020.06.09.141358

**Authors:** Shaoming Sang, Ting Qian, Fang Cai, Hongyan Qiu, Yangqi Xu, Yun Zhang, Qing Zhang, Shajin Huang, Donglang Jiang, Yun Wu, Haiyang Tong, Xiaoli Pan, Changpeng Wang, Xiaoqin Cheng, Kai Zhong, Yihui Guan, Michael X. Zhu, Xiang Yu, Weihong Song, Chunjiu Zhong

## Abstract

**Background:** Thiamine diphosphate (TDP) reduction plays an important role in cerebral glucose hypometabolism, the neurodegenerative indicator, in Alzheimer’s disease (AD). The mechanism underlying TDP reduction remains elusive. Thus, it is critical to define the mechanism and its effect on neurodegeneration, the pathological basis of the disease occurrence and progression.

**Methods:** The mRNA levels of all known genes associated with thiamine metabolism, including *thiamine pyrophosphokinase* (*TPK*), *Solute Carrier Family 19 Member 2 (SLC19A2)*, *SLC19A3*, and *SLC25A19*, in brain samples of patients with AD and other neurodegenerative disorders in multiple independent datasets were analyzed. TPK protein levels were further examined in the brain tissues of AD patients and control subjects. A mouse model with conditional knockout (cKO) of *TPK* gene in the excitatory neurons of adult brain was established.

**Results:** The brain *TPK* mRNA level was markedly lower in AD patients, but not in other neurodegenerative disorders. The brain TPK protein level was also significantly decreased in AD patients. *TPK* gene knockout in the mice caused cerebral glucose hypometabolism, β-amyloid deposition, Tau hyperphosphorylation, neuroinflammation, and neuronal loss and brain atrophy. Cross-species correlation analysis revealed the similar changes of gene profiling between the cKO mice and AD patients.

**Conclusions:** The deficiency of brain TPK, a key enzyme for TDP synthesis, is specific to AD. The cKO mice show AD-associated phenotypes and could serve as a new mouse model for AD studies. Our study provides a novel insight into the critical role of TPK in AD pathogenesis and its potential for the disease treatment.

---

Alzheimer’s disease (AD) is a devastating neurodegenerative disease clinically characterized by progressive memory loss and cognitive decline. The major neuropathological features of AD include β-amyloid (Aβ) deposition, Tau hyperphosphorylation, glial activation, neuroinflammation, glucose hypometabolism, and neurodegeneration characterized by progressive neuronal loss and brain atrophy. Among them, neurodegeneration is the key pathophysiological feature and the direct pathogenic basis for AD occurrence and progression. Aβ deposition has been considered as an initial pathogenic factor of the disease ^1, 2^. However, transgenic rodent models with significant long-term Aβ deposition do not exhibit significant neuronal loss, brain atrophy, or cerebral glucose hypometabolism ^3^, suggesting that Aβ deposition might not be a key pathogenic factor contributing to the disease development. The recent failures of a series of Phase 3 clinical trials targeting Aβ deposition by various approaches such as vaccines ^4^, antibodies ^5–7^, and inhibitors of β and γ-secretases ^8, 9^ reinforces the doubts about Aβ’s pathogenic role in AD. Thus, the relationship between the multiple pathophysiological features and the pathogenesis of AD remains to be clarified.

Brain glucose hypometabolism, the indicator of neurodegeneration, has been considered as the consequence of cerebral dysfunction. However, it has been demonstrated that cerebral glucose hypometabolism is a preclinical characteristic that occurs decades prior to clinical manifestations of the disease ^10, 11^ and it is difficult to fully explain the alteration as a result of brain dysfunction. Furthermore, the activities of three rate-limiting enzymes of glucose metabolism, transketolase in the non-oxidative branch of pentose phosphate pathway, pyruvate dehydrogenase and α-ketoglutarate dehydrogenase in the Krebs cycle, are significantly reduced in AD ^12, 13^. As the critical cofactor of these three enzymes, thiamine diphosphate (TDP), the bioactive form of thiamine, is also markedly decreased in AD but not in other neurological disorders, including Parkinson’s disease, vascular dementia, and frontotemporal lobar degeneration ^14, 15^. Our previous study has demonstrated that TDP reduction is closely correlated with cerebral glucose hypometabolism in AD, whereas Aβ deposition is not ^3^. Thus, to identify the key contributor of TDP reduction in AD may help to understand the disease pathogenesis and provide a novel avenue for AD drug development.

## Methods

The mRNA levels of all known genes associated with thiamine metabolism, including *thiamine pyrophosphokinase* (*TPK*, a key enzyme converting thiamine to TDP) and the three transporters *Solute Carrier Family 19 Member 2* (*SLC19A2*), *SLC19A3*, and *SLC25A19* in brain samples of patients with AD and other neurodegenerative disorders in multiple independent public datasets were analyzed. TPK protein levels in the brain tissues of AD patients and control subjects were further examined using Western blot analysis. A conditional knockout (cKO) mouse model was established, in which the *TPK* gene was specifically disrupted in the glutamatergic neurons of cerebral cortex and hippocampus under the *CaMKII-Cre*^*ERT2/+*^ promoter (Supplementary Figure 1, Figure 2A). Morphological studies with immunofluorescent staining and structural T2-weighted magnetic resonance imaging of mouse brains were performed. Behavior tests, positron emission tomography/computer tomography with 18F-fluorodeoxyglucose (FDG PET/CT), untargeted metabolomics, and RNA-seq analyses were applied (details in Supplementary Methods and Materials).

## Results

### Reduced *TPK* expression in the brain of AD patients

In fusiform gyrus samples of AD patients in the Illumina HiSeq 2500 RNA-seq dataset (GSE95587), *TPK* mRNA levels were significantly reduced as compared with that in the control subjects, while there were no changes in the mRNA levels of the three transporters related to thiamine metabolism (Figure 1A, Supplementary Table 1). *TPK* mRNA levels were found to have a negative correlation with Braak staging scores in AD patients (R = 0.31, P = 0.079, Figure 1B). In addition, for subjects with Braak stage IV pathology, control samples exhibit significantly higher TPK mRNA levels than AD samples (Figure 1C).

**Figure 1.**
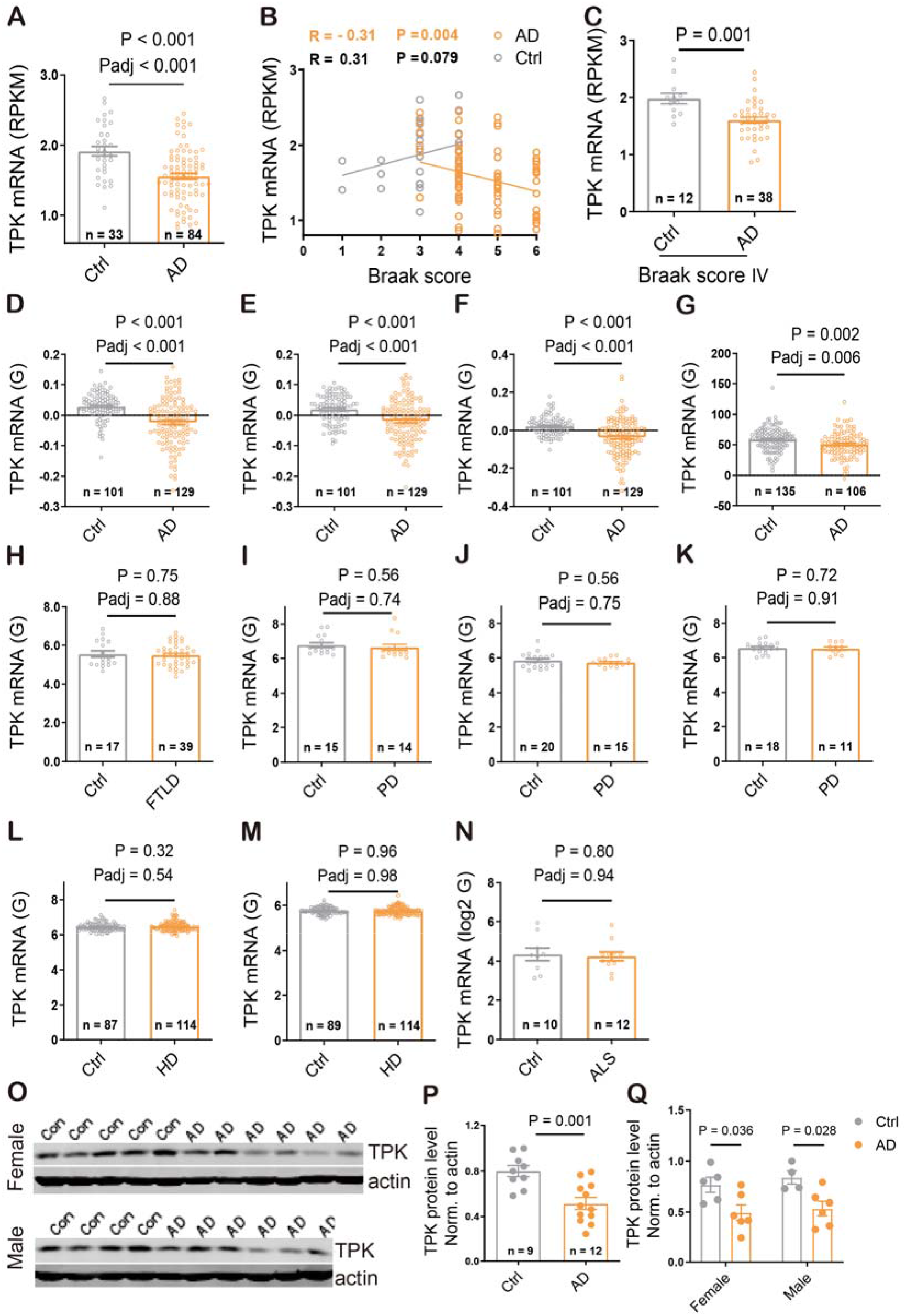
Specific down-regulation of TPK expression in brain samples of AD patients. A-C. Expression analysis of *TPK* mRNA using the GSE95587 dataset measuring fusiform gyrus cortex samples of AD. A. *TPK* mRNA levels in brain samples of AD patients were significantly lower than in that of control subjects. B. Correlation between brain *TPK* mRNA levels and scores of Braak staging: while the correlation was negative for AD patients (n = 84), it tended to be positive for control subjects (n = 33). C. For individuals with Braak stage IV pathology, control subjects showed significantly higher levels of *TPK* mRNA than AD patients. D-G. The levels of TPK mRNA were significantly reduced in brain samples of AD patients as compared with that in control subjects in the GSE15222 dataset measuring frontal cortex samples of AD (D), GSE44768 dataset measuring cerebellar samples of AD (E), GSE44770 dataset measuring dorsolateral prefrontal cortex samples of AD (F), and GSE44771 dataset measuring visual cortex samples of AD (G). H. There is no significant difference in TPK mRNA levels between patients with frontotemporal lobar degeneration and control subjects. I-K. The levels of TPK mRNA show no significant differences in brain samples of Parkinson’s disease (PD) patients as compared with that in control subjects in the GSE20168 dataset measuring prefrontal cortex samples of PD (I), GSE20291 dataset measuring putamen samples of PD (J), and GSE20292 dataset measuring substantia nigra samples of PD (K). L & M. The levels of TPK mRNA in brain samples of Huntingdon’s disease (HD) patients exhibit no significant differences as compared with that in control subjects in the GSE3790A dataset (L) and GSE3790B dataset measuring cerebellum, frontal cortex and caudate nucleus samples of HD (M). N. There was no significant difference in TPK mRNA levels between patients with amyotrophic lateral sclerosis and control subjects. O. Images of western blot of TPK protein in the cortical tissues of AD and control subjects. P. Quantification of TPK protein levels in brain samples of AD and control subjects. The results showed significant reduction of TPK protein level in brain samples of AD patients compared to control subjects (P = 0.001). Q. The levels of TPK protein in brain samples of both male and female AD patients were also significantly decreased as compared with that in respectively control subjects (Male: n = 4, 6 for control and AD subjects, P = 0.036; Female: n = 5, 6 for control and AD subjects, P = 0.028; P = 0.999 for male vs female AD cases; P = 0.990 for male vs female control subjects).

The reduction of *TPK* mRNA levels in AD patients was verified in other open-source datasets, including GSE15222, GSE44768, GSE44770, and GSE44771 (Figure 1D-G), while the mRNA levels of thiamine transporters had either no significant changes or slight increase (Supplementary Table 1). There were no significant changes in the mRNA levels of *TPK*, *SLC25A19*, *SLC19A2*, and *SLC19A3* in samples of frontotemporal lobar degeneration, Parkinson disease, Huntington’s disease, and amyotrophic lateral sclerosis (Figure 1H-N and Supplementary Table 1).

Next, we examined TPK protein in the brain samples. Cortical tissues of the frontal lobe from 12 AD patients and 9 control subjects were homogenized and analyzed by Western blot. There was no significant difference in the average age of the AD patients (70.83 ± 2.74 years) and control subjects (65.44 ± 4.41 years) (P = 0.2899). TPK protein levels were also significantly decreased in the cortical tissues of AD patients compared to the control subjects, (P = 0.001, Figure 1O, P). Furthermore, both male and female AD patients showed significant reduction of TPK protein level and there was no difference between male and female patients (Figure 1O, Q).

### Dysfunction of brain glucose metabolism in the *TPK* knockout mice

A mouse model with conditional *TPK* knockout in the excitatory neurons of adult brain was generated (Figure 2A). The levels of TPK protein and TDP in the mouse brains were measured at 10 weeks after tamoxifen treatment, and both TPK and TDP levels in the cKO mouse brains were significantly reduced as compared with those of control littermates (*Tpk*^*fl/fl*^, Figure 2B, C). Blood TDP levels were not significantly affected in the cKO mice (Supplementary Figure 2A). The body weight was comparable between the cKO and control mice at all observation time points except in the 8^th^ week after tamoxifen treatment, at which the body weight was significantly increased in the cKO mice than the control littermates (Supplementary Figure 2B).

**Figure 2.**
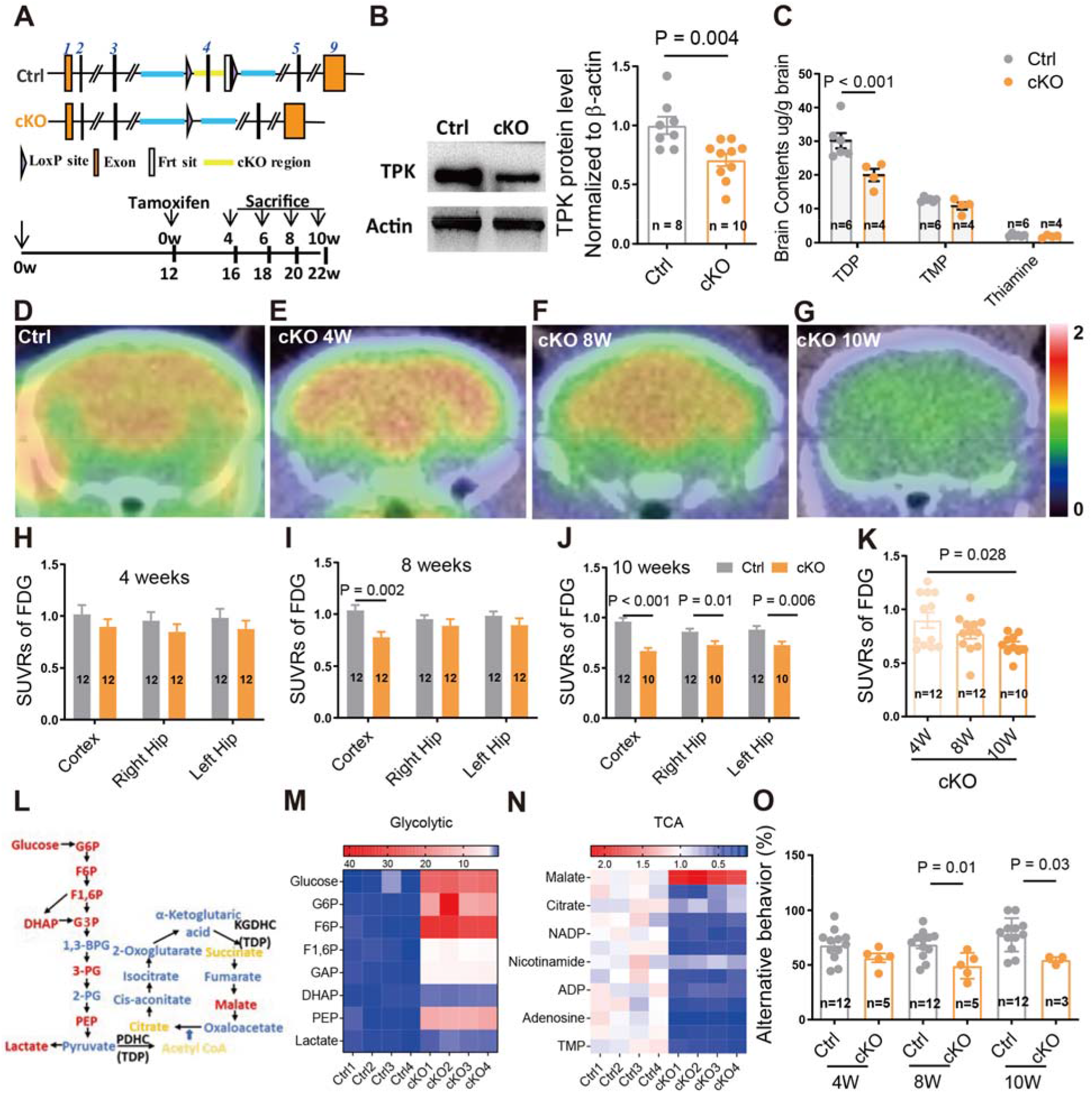
Altered brain glucose metabolism in *TPK* conditional knockout mice. A. Strategy used to generate adult cKO mice with conditional knockout of *TPK* in brain excitatory neurons. Tamoxifen was given for five consecutive days when animals were 12 weeks old. B. Confirmation of a significant reduction of TPK protein in brain samples of the cKO mice as compared to control littermates. C. The levels of TDP, but not that of thiamine monophosphate (TMP) and thiamine, in brain samples of the cKO mice were significantly decreased as compared to that of the control littermates. For both f and g, samples were from 10^th^ week after tamoxifen induction. D-G. Representative brain images of FDG PET/CT of the control littermates (D) and the cKO mice in the 4^th^ (E), 8^th^ (F), and 10^th^ (G) week after tamoxifen treatment. H-J. Statistics on the levels of cortical and hippocampal FDG uptake between the control littermates and the cKO mice in the 4^th^ (H), 8^th^ (I), and 10^th^ (J) week after tamoxifen treatment. No difference was found at the 4^th^ week. In the 8^th^ week, there was a significant decline in cortical but not hippocampal FDG uptake. In the 10^th^ week, there were significant declines in both cortical and hippocampal FDG uptake. K. The levels of cortical FDG uptake in the cKO mice manifested progressive declines in the 4^th^, 8^th^, and 10^th^ week after the tamoxifen treatment. L. Schematic representation of glycolysis and oxidative phosphorylation pathways, with red, yellow, and blue indicating increase, decrease, and no alternation, respectively, in the cortices of the cKO mice. M & N. Heatmaps of quantification of metabolic intermediates of glycolysis: glucose, glucose-6-phosphate, fructose-1,6-phosphate, lactate, phosphoenolpyruvate (M), and oxidative phosphorylation (TCA) pathway: succinate, citrate, Acetyl CoA, ATP, NAD, and NADP, malate (N). O. Alternation of Y-maze test of the control littermates and the cKO mice in the 4^th^, 8^th^, and 10^th^ week after tamoxifen treatment. Significant decreases were found in the 8^th^, and 10^th^ week, but not the 4^th^ week. Summary data represent means ± SEM for numbers of tests or subjects indicated in parentheses. P values are indicated above the bars or > 0.05 if not labeled.

Compared to the control littermates, the cKO mice exhibited a significant decrease in glucose uptake initially in the cortex in the 8^th^ week, and then extended to most brain regions in the 10^th^ week after tamoxifen treatment (Figure 2D-J, Supplementary Figure 2C-E). Self-comparison analysis showed a gradual decline of glucose uptake in the cKO cortex (Figure 2K). Moreover, metabolic analysis of the cortex of the cKO mice in the 10^th^ week after tamoxifen treatment revealed significant accumulation of many substrates involved in glycolysis but decreased levels of those involved in oxidative phosphorylation (Figure 2L-O). Y-maze assay showed significant decreases in spatial working memory of the cKO mice in the 8^th^ week after tamoxifen treatment (Figure 2O). Unfortunately, the Morris water maze was difficult to be analyzed due to the increased body weight of the cKO mice and their reluctance to move (Supplementary Figure 3).

### Progressive neuronal loss and brain atrophy in the *TPK* cKO mice

There were no significant changes in the brain volumes of the cKO mice for the first 6 weeks after tamoxifen treatment, after which a progressive decline was detected by longitudinal magnetic resonance imaging (Figure 3A, B). This was supported by the measurements of brain superficial areas, total brain weights, cortical weights, and hippocampal weights (Figure 3C, D; Supplementary Figure 4A-C). No changes were found in the cerebellum weight of the cKO mice except for a slight decrease in the 10^th^ week (Supplementary Figure 4D). Histological examination further demonstrated a progressive loss of cerebral neurons in the cKO mice beginning in the 8^th^ week after tamoxifen treatment (Figure 3E-J, Supplementary Figure 4E, F). The loss of cortical synapses in the cKO mice began earlier than the neuronal loss and was detectable in the 6^th^ week (Supplementary Figure 4G-J). In addition, the timing of neuronal loss and brain atrophy closely matched with the onset of cognitive impairment in the cKO mice shown above.

**Figure 3.**
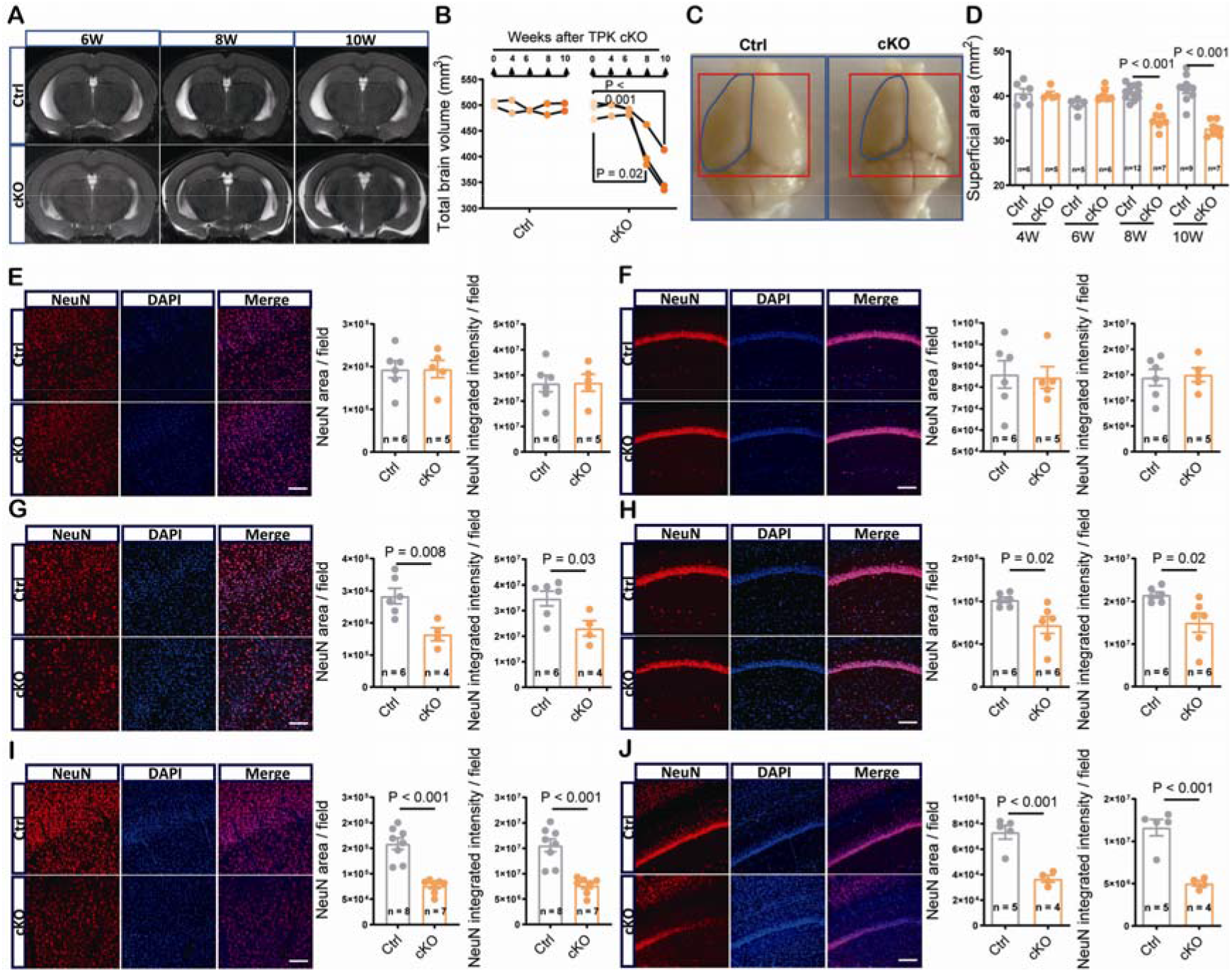
*TPK* cKO mice manifest progressive neuronal loss and brain atrophy. A. Representative images of structural brain T2 weight magnetic resonance imaging (MRI) of the control and cKO mice in the 6^th^, 8^th^, and 10^th^ week after tamoxifen treatment. B. Quantification of total brain volumes based on MRI showing progressive declines of the cKO mice (n = 3) in the 8^th^ and 10^th^, but not 4^th^ and 6^th^ week after tamoxifen treatment as compared with the self-brain volumes before tamoxifen treatment. The brain volumes of the control littermates had no significant changes at all observation time points (n = 2). C. Representative brain samples of the control littermates and cKO mice in the 10^th^ week after tamoxifen treatment. D. Quantification of brain superficial areas showing significant reductions in the cKO mice in the 8^th^ and 10^th^, but not 4^th^ and 6^th^, week, as compared to the control littermates. E-J. Representative images (*left*) and quantification of areas (*middle*) and integrated intensities (*right*) of NeuN-positive cells in cortical (E, G, I) and hippocampal (F, H, J) regions of the control littermates and cKO mice in the 6^th^ (E & F), 8^th^ (G & H), and 10^th^ (I & J) week after tamoxifen treatment. For both cortex and hippocampus, significant decreases were found in the cKO mice as compared to the control littermates in the 8^th^ and 10^th^ week, but not in the 6^th^ week, when quantified by either area or intensity. Summary data represent means ± SEM for numbers of tests indicated in parentheses. Scare bar, 100μm. P values are indicated above the bars or > 0.05 if not labeled.

### Aβ deposition, Tau hyperphosphorylation, and neuroinflammation in the *TPK* cKO mice

Aβ deposition, Tau hyperphosphorylation and neuroinflammation are the neuropathological hallmarks of AD brains. Cortical samples of the cKO mice in the 10^th^ week after tamoxifen induction showed significant increases in both soluble and insoluble, as well as total, Aβ42 and Aβ40 levels as compared to the control littermates. The ratios of total Aβ42 to total Aβ40 and insoluble Aβ42 to insoluble Aβ40 in the cKO mice were also increased as compared to the controls while the ratio of soluble Aβ42 to soluble Aβ40 was not affected (Figure 4A-C). Furthermore, the levels of Aβ precursor protein (APP) and β-site APP cleaving enzyme 1 (BACE1) were significantly elevated in cortices of the cKO mice. The cKO mice also had increased APP expressions, but not BACE1 expressions in the hippocampi as compared to the control littermates (Figure 4D-I).

**Figure 4.**
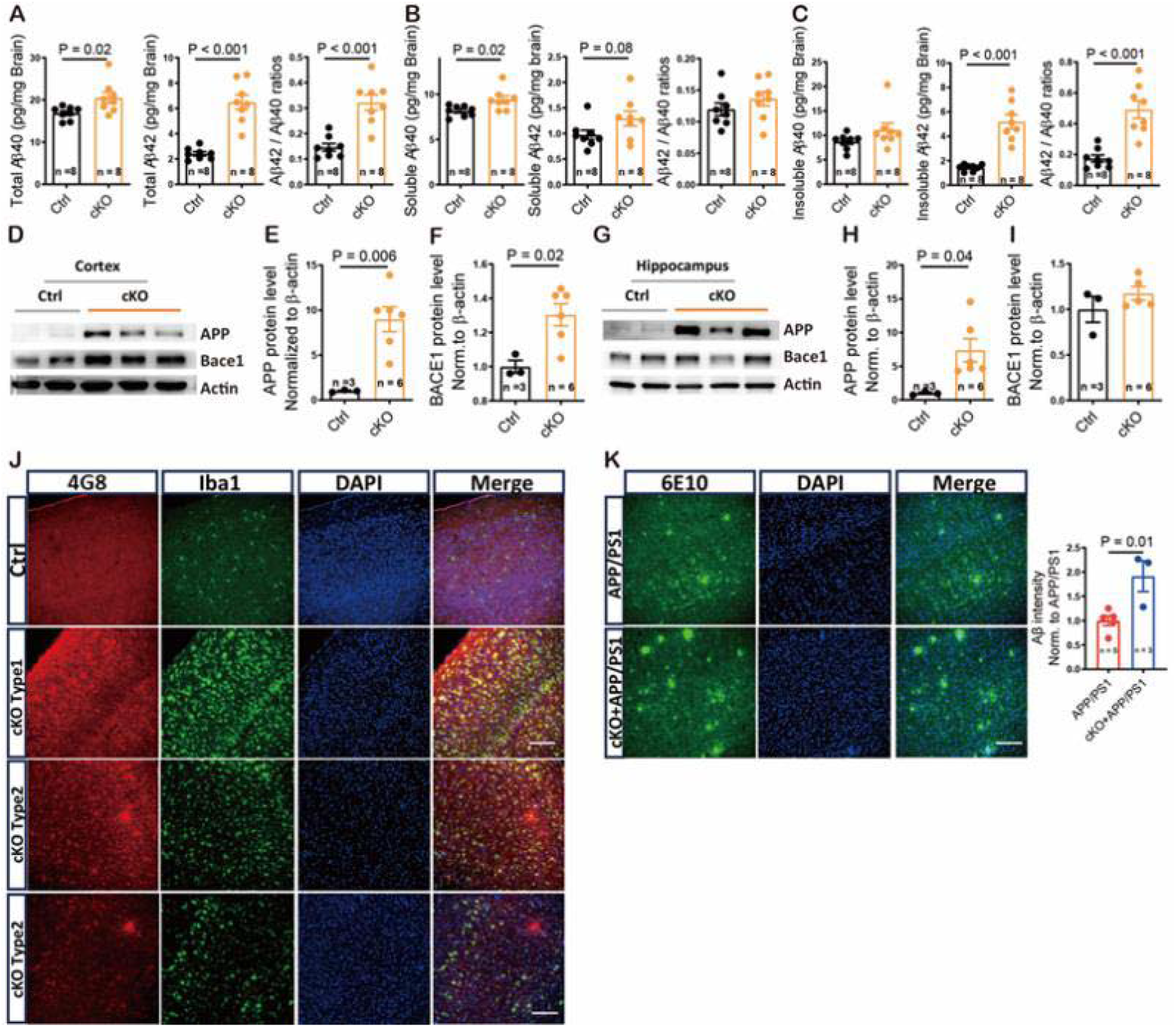
Brain Aβ deposition in *TPK* cKO mice. A-C. Comparisons of levels of total (A), soluble (B) and insoluble (C) Aβ40, Aβ42, and Aβ42/Aβ40 ratios in brain samples between the control littermates and cKO mice in the 10^th^ week after tamoxifen treatment. Significant increases were found in total Aβ40, total Aβ42, total Aβ42/total Aβ40 ratio, soluble Aβ40, insoluble Aβ42 and insoluble Aβ42/insoluble Aβ40 ratio. D-I. Comparisons of APP and BACE1 protein levels in cortical (D-F) and hippocampal (G-I) samples of the control littermates and cKO mice in the 10^th^ week after tamoxifen treatment by Western blot. Representative blots (D & G) and quantifications (E, F, H, I) are shown. In the cortex, both APP and BACE1 levels were significantly increased, while in the hippocampus, only APP, but not BACE1 protein, levels were significantly increased in the cKO mice. J. Representative images of immunofluorescent staining show that Aβ enhancement is mainly in microglia without plaque formation in the brain regions with significant microglial activation (we defined this phenomenon as type I Aβ deposition) or appears as plaques in the brain areas with mild to moderate microglial activation (we defined this phenomenon as type II Aβ deposition) in the 10^th^ week after tamoxifen treatment. K. Aβ plaques detected by immunofluorescent staining in the cortex of the APP/PS1 littermates and cKO/APP/PS1 mice in the 10^th^ week after tamoxifen treatment. Representative images (left) and quantification of Aβ plaques (right) are shown. Aβ plaques were significantly increased in the cortex of the cKO/APP/PS1 mice as compared to the APP/PS1 mice. Scare bar, 100μm. Summary data represent means ± SEM for numbers of tests indicated in parentheses. P values are indicated above the bars or > 0.05 if not labeled.

Immunofluorescent staining showed that cortical Aβ enhancement was mainly observed two types: In brain regions with significant microglial activation, Aβ existed mainly in the microglia but not the neurons (we defined as type I); In brain regions with mild or moderate microglial activation, plaques were found (we defined as type II. Figure 4J and Supplementary Figure 5A-B). Immunofluorescent staining showed that TPK protein was mainly enriched in neurons but less in microglia in the brains of wild-type mice, while our conditional knockout of the *TPK* gene in the excitatory neurons of adult brain resulted in abolishing the TPK protein expression in neurons but not in microglia of the cKO mice (Supplementary Figure 5 C-D). To further examine the effect of TPK on plaque formation, the APP/PS1 transgenic mice were cross-bred with the cKO mice to generate the APP/PS1/cKO mice. We found that the plaque formation was significantly enhanced in the cortices of the APP/PS1/cKO mice as compared with the APP/PS1 transgenic mice in both the 6^th^ and 10^th^ week after tamoxifen treatment (Figure 4K, Supplementary Figure 5E). The data demonstrated that TPK deficiency facilitates Aβ plaque formation *in vivo*.

Significant enhancement of Tau phosphorylation was observed in both cortical and hippocampal tissues of the cKO mice beginning in the 8^th^ week after tamoxifen treatment as compared to the control littermates (Figure 5A, B, Supplementary Figure 6A-F). Silver staining and Immunofluorescent staining with AT8 antibody detected significant tangles in neurons in the cKO mouse brains (Supplementary Figure 6G-I).

**Figure 5.**
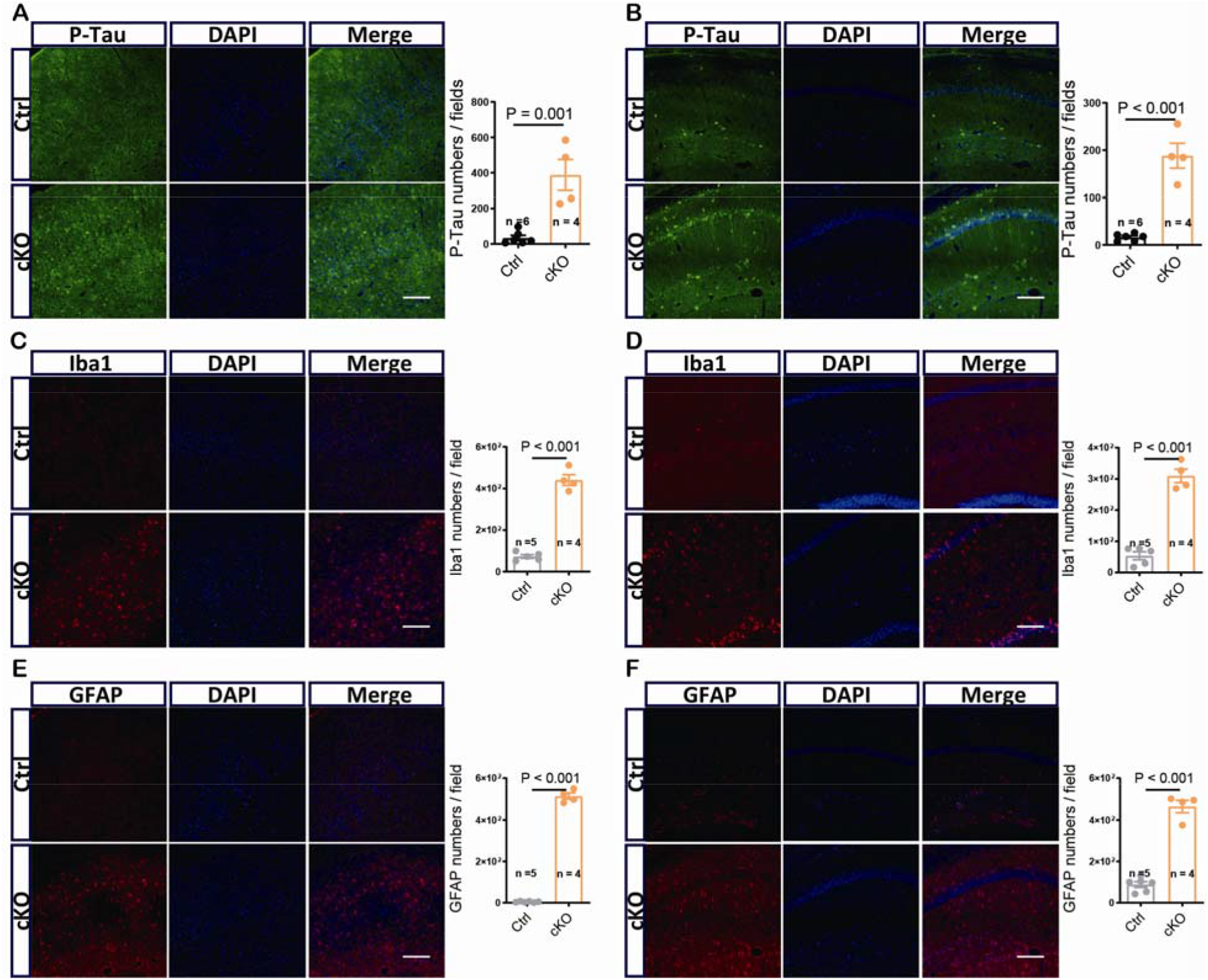
Brain Tau hyperphosphorylation, and neuroinflammation in *TPK* cKO mice. A & B. Tau phosphorylation in the cortex (A) and hippocampus (B) of the control littermates and cKO mice in the 10^th^ week after tamoxifen treatment, as revealed by immunofluorescence. Representative images (left) and quantification of phosphorylated Tau (right) are shown. Tau phosphorylation was significantly increased in both the cortex and hippocampus of the cKO mice compared to the control littermates. C & D. Cortical (C) and hippocampal (D) microglia in the control littermates and cKO mice in the 10^th^ week after tamoxifen treatment, as revealed by immunofluorescence for Iba1. Shown are representative images (left) and quantification (right) for the density of Iba1+ cells. E & F. Similar to C and D, but for astrocytes revealed by immunofluorescence for GAFP. Both cortex and hippocampus showed significant increases in the numbers of microglia and astrocytes in the cKO mice as compared to the control littermates. Scare bar, 100μm. Summary data represent means ± SEM for numbers of tests indicated in parentheses. P values are indicated above the bars or > 0.05 if not labeled.

The cKO mice showed significant activation of microglia and astrocytes in cortical tissues beginning in the 4^th^ week after tamoxifen treatment. In the hippocampus, the activation of microglia and astrocytes started from the 6^th^ and 8^th^ week, respectively, which was later than that in the cortex (Figure 5C-F, Supplementary Figure 6). The levels of tumor necrosis factor-α (TNF-α) in the cKO brains were significantly enhanced starting from the 4^th^ week after tamoxifen treatment, while the levels of other factors, such as interleukin-6 and interleukin-1β, were increased starting from the 6^th^ week (Supplementary Figure 7).

### Gene profiling revealed similar changes in the *TPK* cKO mice and AD patients

RNA sequencing of cortical samples in the 10^th^ week after tamoxifen treatment showed that over 4,000 genes were significantly upregulated and over 4,000 genes were downregulated in the cKO mice as compared to the control littermates (Figure 6A). Based on gene ontology (GO), gene set enrichment analysis (GSEA), and the Kyoto Encyclopedia of Genes and Genomes (KEGG) pathway analysis, we found that genes associated with multiple death signal pathways and neuroinflammation including cellular senescence, apoptosis, p53 signaling pathway, cytosolic DNA-sensing pathway, ferroptosis, and necroptosis, were up-regulated in the cKO cortex, and these alterations may contribute to the neuronal and synaptic loss (Figure 6C). Cytokine-cytokine receptor interaction, and nucleotide-binding oligomerization domain (NOD)-like receptor, Toll-like receptor and TNF signaling pathways were also upregulated, which may underlie glial activation and enhanced expression of inflammatory factors (Figure 6C). Down-regulated were the genes strongly involved in neurotransmitter pathways, including glutamatergic synapse, neuroactive ligand-receptor interaction, synaptic vesicle cycle, and long-term potentiation (Figure 6B).

**Figure 6.**
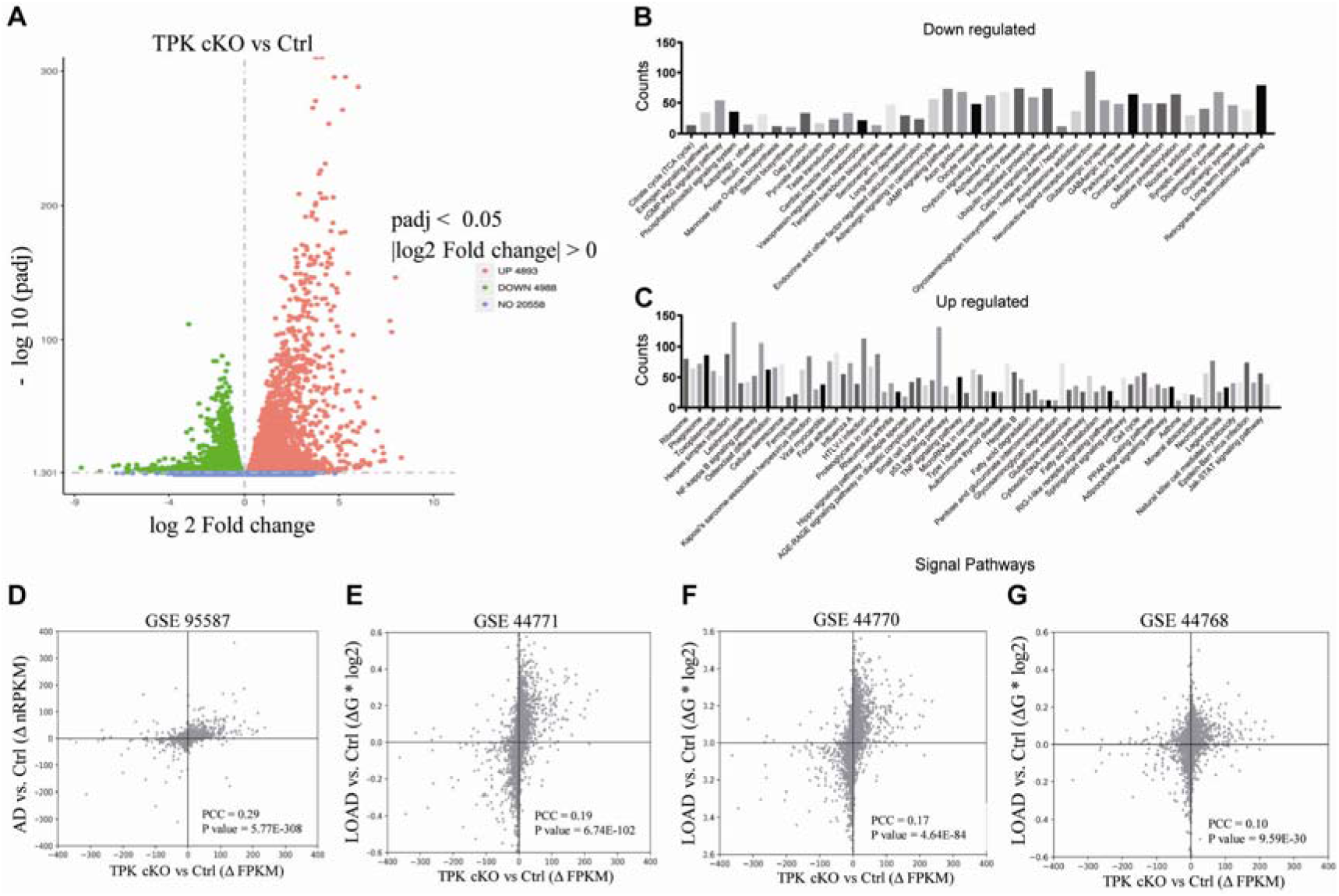
Genome-wide expression changes in *Tpk* cKO mice closely correlate with those in AD patients. A. Transcriptional profiles of the cKO mouse cortex in the 10^th^ week after tamoxifen induction. B. Down-regulated genes in the cKO mice are enriched of those involved in neurotransmission pathways. C. Up-regulated genes in the cKO mice are enriched of those involved in cell death pathways. D-G. Scatter plots showing genome-wide correlated changes in gene expression between the cortex samples of the cKO mice and brain samples of AD patients (D: fusiform gyrus, PCC = 0.29, P = 5.77e-308; E: prefrontal cortex, PCC = 0.19, P = 6.74e-102; F: visual cortex, PCC = 0.17, P = 4.64e-84; G: cerebellar cortex: ACC = 0.10, P = 9.59e-30).

Changes in genome-wide transcription levels were compared between the cKO mice and AD patients by cross-species correlation analysis. The results showed that the changes found in the cKO mice were highly correlated with that in the fusiform gyrus (GSE95587), prefrontal cortex (GSE44771), and visual cortex (GSE44770) of AD patients (Figure 6D-F, Supplementary Table 2), but not strongly correlated with that in cerebellar cortex of AD (GSE44768), or in the samples of frontotemporal lobar degeneration (GSE13162), Parkinson’s disease (GSE20295), Huntingdon’s disease (GSE3790), or amyotrophic lateral sclerosis (GSE18920) (Figure 6G, Supplementary Figure 8 and Supplementary Table 2). To further identify the genes and pathways that are commonly regulated both in the cKO mice and AD patients, we further performed GSEA on AD datasets of GSE99587. Among the top 20 modules of significantly decreased transcriptions, 12/20 modules were enriched in oxidative phosphorylation and mitochondria energy metabolism, another 8/20 modules were enriched in synaptic/neuronal function and neurotransmission. Among the top 20 modules of significantly increased transcriptions, 9/20 were enriched in astrocyte/microglia activation and chemokine secretion pathways, 2/20 were enriched in apoptosis pathway (Supplementary Figure 9), and others in long chain fatty acid biosynthetic, small ribosomal subunit, etc. The changes in those modules were further confirmed by a recent large-scale proteomic study. Importantly, the glucose metabolism module was also identified as having the strongest correlations with AD pathological features ^16^. These analyses further support a causative effect of TPK deficiency in AD.

## Discussion

Here, we identified that *TPK*, the only thiamine metabolism related gene, was specifically downregulated in the brain of AD patients but not of patients with other neurodegenerative diseases, including frontotemporal dementia, Parkinson’s disease, Huntington’s disease, and amyotrophic lateral sclerosis (Figure 1). Furthermore, increasing TPK expression may be protective against AD, as indicated by the positive correlation between TPK mRNA levels and the scores of Braak staging in control subjects (R = 0.31, P = 0.079) but by the negative correlation in AD patients (R = −0.31, P = 0.004, Figure 1B), and the significantly higher TPK mRNA levels in control subjects than AD patients with the pathological feature of Braak stage LJ (Figure 1C). Importantly, the TPK protein level was also significantly decreased in the cortical tissues of AD patients compared to control subjects. These results indicate that impairment in TPK expression/function may be a causative factor contributing to AD pathogenesis.

The studies of the cKO mice further confirmed the aforementioned results and supported a causative relationship between TPK deficiency and the pathophysiological features of AD, including progressive neuronal loss and brain atrophy (Figure 3), Aβ deposition (Figure 4), Tau hyperphosphorylation, glial activation and neuroinflammation (Figure 5), and glucose hypometabolism (Figure 2, Supplementary Figure 2). Our previous study has shown that inhibiting the expression of *TPK*, *SLC25A19*, or *SLC19A3* in individual neurons with RNA interference significantly reduced dendritic complexity and soma size ^17^, suggesting a coordinated regulation of soma and dendrite growth by thiamine metabolism. It is reasonable to consider that TDP reduction caused by TPK deficiency, rather than the loss of TPK’s own function *per se*, serves as a pathogenic factor contributing to AD pathogenesis. Therefore, having demonstrated that the cKO mice displayed neuronal loss and brain atrophy, as well as many other pathophysiological features of AD, our study strongly supports that TPK deficiency and the consequent impairment in thiamine metabolism in the brain constitute a pathogenic factor that underlies multiple pathophysiological features of AD and plays a critical role in disease pathogenesis. Ours and another previous study have shown that TPK activity was not significantly changed in brain and blood samples of AD patients compared to control subjects ^18,19^. It is possible that the existing method of TPK activity measurement was insensitive due to the condition of optimal pH and Mg2+ concentration, as well as abundant substrates (ATP and thiamine). More sensitive methods for evaluating TPK activity should be established in the future study.

Aβ deposition is undoubtedly a pathological hallmark of AD. However, the mechanism of Aβ deposition in sporadic cases that account for the majority of AD cases remains to be elucidated. Our present study demonstrates that TPK deficiency in excitatory neurons promotes Aβ deposition by enhancing the expression of APP and BACE1 proteins in the mouse brain, indicating that alterations in thiamine and glucose metabolism precede and cause brain Aβ deposition, in contrast to the hypothesis derived from studying the disease biomarker dynamics based on familial AD, which suggests that Aβ deposition is an initial factor ^20^. Interestingly, Immunofluorescent analysis showed that Aβ enhancement was mainly occurred in microglia in brain regions with significant microglial activation but not in neurons and there was also plaque formation in brain regions with mild to moderate microglial activation in the cKO mice. This was verified by further study on the APP/PS1/cKO mice carrying human mutant genes. The TPK deficiency enhanced significantly cortical Aβ plaques of the APP/PS1/cKO mice (Figure 4J, K, Supplementary Figure 5E).

Neuroinflammation is another important pathophysiological feature of AD. Our study showed that glial activation and the increase in TNF-α occurred at the early stage (the first month) of TPK deficiency when the cKO mice exhibited no signs of neuronal loss. Thus, the abnormal intracellular processes of glucose metabolism due to TPK deficiency caused the glial activation and neuroinflammation, indicating that the brain glucose hypometabolism could be the inducer of neuroinflammation in the brain of AD patients. Future studies should address how inhibition of TPK in neurons triggers glial activation and neuroinflammation and what roles the early glial activation and neuroinflammation play in neuronal loss and other pathophysiological features associated with TPK deficiency.

To date, no animal models can fully recapitulate all pathophysiological features of AD and their causal relationships. The lack of a proper animal model may be one of the major reasons for the failures of the phase 3 clinical trials on new drugs aiming at preventing ^21, 22^ and/or treating AD ^4–9^. The current animal models either possess only a single AD-related pathophysiological characteristic (e.g., Aβ deposition) or multiple (usually two) but independent characteristics (e.g., Aβ deposition and Tau hyperphosphorylation in APP/PS1/Tau transgenic mice). These models do not represent the human disease; nor do they reveal the intrinsic relationships among the multiple pathophysiological features of AD. The *TPK* cKO mice described here represent an excellent AD animal model: first, TPK deficiency and the consequent thiamine and glucose hypometabolism are major pathophysiological features of AD; second, TPK deficiency in excitatory neurons of adult mice resulted in a comprehensive manifestation of nearly all other pathophysiological features typically found in AD patients; third, transcriptome analysis further indicates that the neuronal *TPK* gene knockout in adult mice triggered pathophysiological responses closely resembling that in AD patients. Therefore, the *TPK* cKO mouse model provides an excellent tool to explore the causal relationships and interactions among many of the pathophysiological features of AD, paving the way for more in-depth understanding of AD pathogenesis. In addition, since knocking out of the *TPK* gene in neurons could be manipulated at any age using the cKO mouse model, it offers an opportunity, in future studies, to investigate the age-dependent neurodegeneration of AD, another important characteristic of the disease not assessable by the existing animal models. It is expected that by comparing differences in pathophysiological features between the conditional *TPK* knockouts at different ages, the age-dependent effects could be revealed.

There is no disease-modifying drug for AD treatment. Given that phase 3 clinical trials for AD drugs have failed and reducing brain Aβ deposition ^3–8^ or inhibiting tau aggregation^23^ has little effect on AD progression, new therapeutic strategies are urgently needed. The mechanism underlying the thiamine deficiency in AD patients is not clear, and simple thiamine supplement treatments do not display beneficial effects for AD^24, 25^. Our current study shows that TPK deficiency in mice causes multiple AD-like pathophysiological features, especially neuronal loss and brain atrophy. These results suggest that the abnormality of thiamine metabolism in AD is not due to the substrate deficiency, but the alteration of its conversion process. The future clinical studies should further explore the relationship between TPK expression and the occurrence and progression of the disease, as well as the possibility of TPK as a therapeutic target of AD.

In summary, for the first time, our study unveils that the inhibition of TPK as a key enzyme of thiamine metabolism is AD-specific. Furthermore, TPK deficiency in mice triggers almost all AD-associated phenotypes, including glucose metabolism dysfunction, Aβ deposition, Tau hyperphosphorylation, glial activation and neuroinflammation, progressive neuronal loss leading to brain atrophy. TPK plays a protective role against AD and, therefore, is a potential therapeutic target for the prevention and treatment of this devastating disease. The efficiency of TPK agonist or activator should be explored in future clinical trials for AD treatment. The cKO mice represent a valuable model to further decipher the relationships among multiple pathophysiological features of the disease. The current results provide novel insights into AD pathogenesis and will inspire new strategies in developing drug therapies for AD.

## Acknowledgements

We thank Dr. Min Jiang and Molecular and Cellular Imaging Facility of Institutes of Brain Science (IOBS), Fudan University, for support with confocal microscopy and data analysis, and Dr. Qian Huang and Animal Behavior Facility of IOBS, Fudan University, for support with animal behavioral experiments.

## Supplementary Materials

Supplementary Materials and Methods

Supplementary Figure 1-9

Supplementary Tables 1-2

