## Supplementary Figures and Tables for "Thiamine pyrophosphokinase deficiency induces Alzheimer’s pathology"

**Supplemental Figures and Tables:**

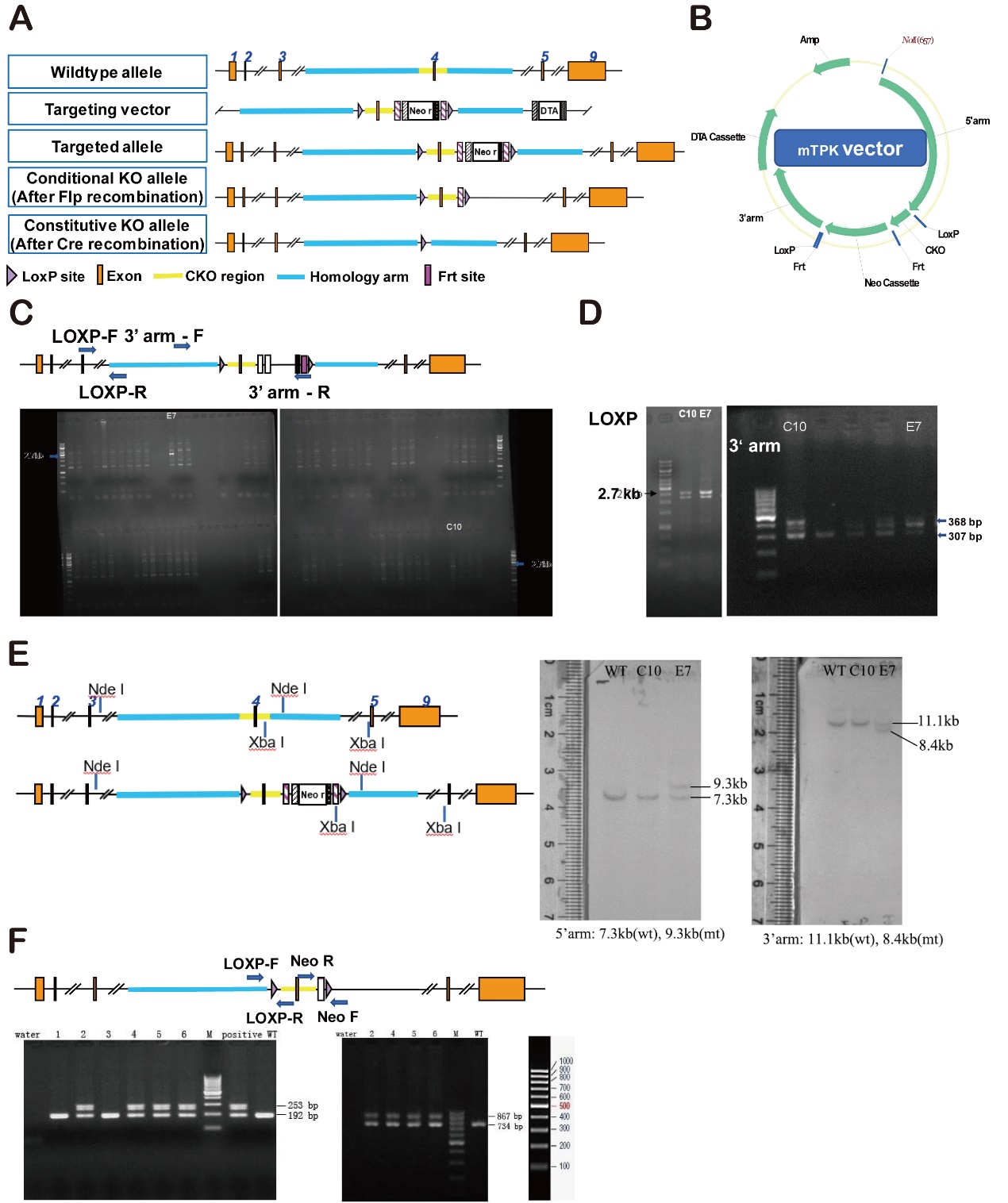

**Figure S1. Establishment of conditional *Tpk* gene knockout mice.** A. Overview of the targeting strategy. Construct of the targeting vector, in which the Neo cassette is flanked by Frt sites and the cKO region flanked by LoxP sites. B. DTA was used for negative selection. C. From 85 clones screened with 3’ arm F / 3’ arm R primers for a ~2700-bp product, 2 positive clones, C10 and E7, were identified. Re-confirmation of the targeted clones with 3’ arm F / 3’ arm R primers showing the ~2700-bp fragment in C10 and E7. D. The clones were further verified with LOXP-F / LOXP-R primers for production of a 307-bp fragment from the wild-type allele and a 368-bp fragment from the recombinant allele. E. Southern Blot analysis conducted by 5’-probe and 3’-probe. For 5’-probe, genomic DNA’s of clones C10 and E7 were digested by *Nde* I and analyzed by Southern blotting for a 7.3-kb band from wild type allele and a 9.3-kb band from recombinant allele; E7 was found positive. For 3’-probe, genomic DNA’s of clones C10 and E7 were digested by *Xba* I and analyzed by Southern blotting for an 11.1-kb band from wild type allele and an 8.4-kb band from recombinant allele; E7 was found positive. F. PCR identification of the F1 mice. Out of 6 pups, 4 (pups 2, 4, 5, 6 from clone E7) were identified positive by the first PCR screening with primers LoxP_F / LoxP_R with the expected 192-bp fragment from wild type allele and 253-bp fragment from recombinant allele (left gel). The tails of the 4 positive pups were recut for the second PCR reconfirmation with a different pair of primers Neo _F / Neo _R and all of them yielded the 867-bp fragment expected from recombinant allele, in addition to the 734-bp fragment from the wild type allele.

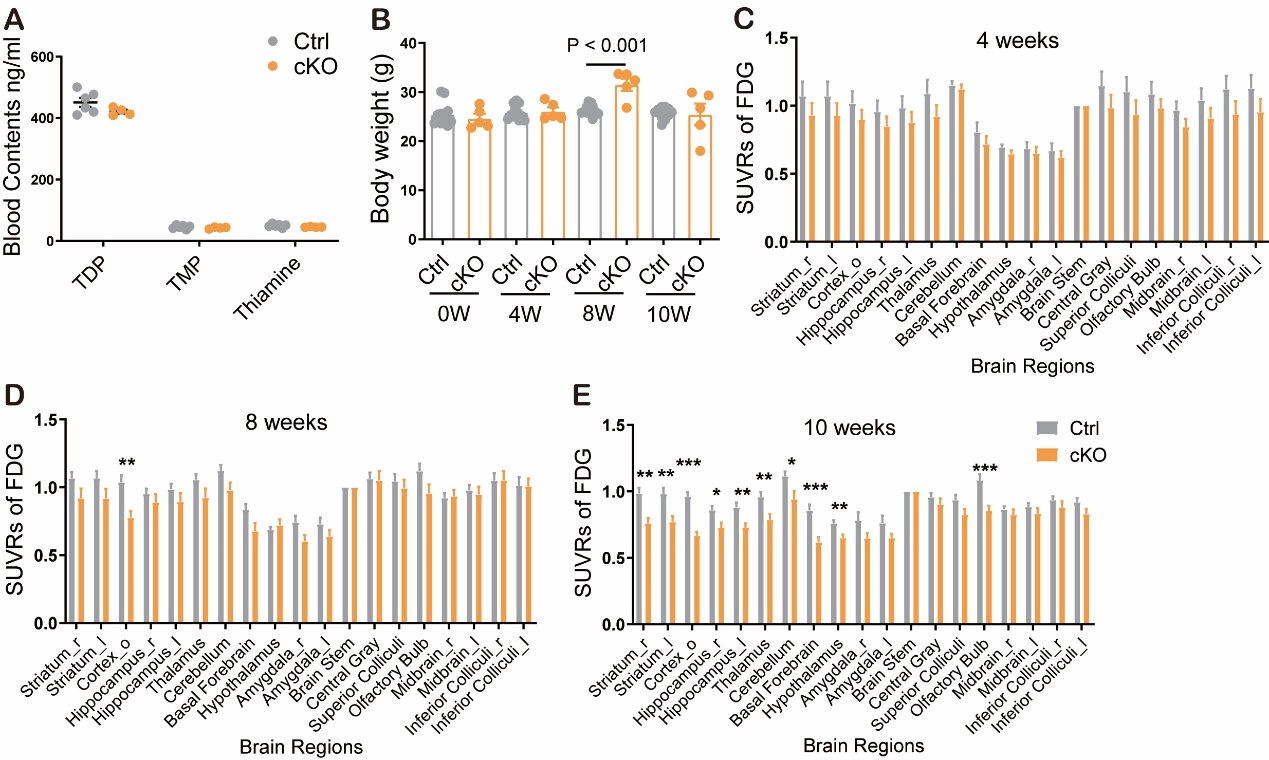

**Figure S2. *Tpk* cKO mice exhibit significant dysfunction of glucose metabolism.** A. There were no significant changes in levels of TDP, thiamine monophosphate (TMP), and thiamine of blood samples in the cKO mice (n = 6) compared to the control littermates (n = 4, P > 0.05). B. The comparison of body weights between the control littermates (n = 12) and cKO mice (n = 5). Except for a significant increase in the cKO mice compared to the control mice at the 8^th^ week (8W, P < 0.001), no differences were found at the 0 week of (0W), 4^th^ (4W), and 10^th^ (10W) weeks (P > 0.05 for all) after tamoxifen treatment. C-E. Levels of FDG uptake in different brain regions of the cKO mice. No significant changes were found at the 4^th^ week in all the brain regions examined (C, n = 12, P > 0.05). A significant decrease appeared in the cortex (P = 0.002), but not in other brain regions of the cKO mice at the 8^th^ week (D, n = 12, P > 0.05). Significant decreases were detected in the cortex, hippocampus, striatum, thalamus, cerebellum, basal forebrain, hypothalamus, and olfactory bulb of the cKO mice (n = 10) compared to the control littermates (n = 11, P < 0.05 for all). Yet, there were still many unaffected areas at the 10^th^ week (E).

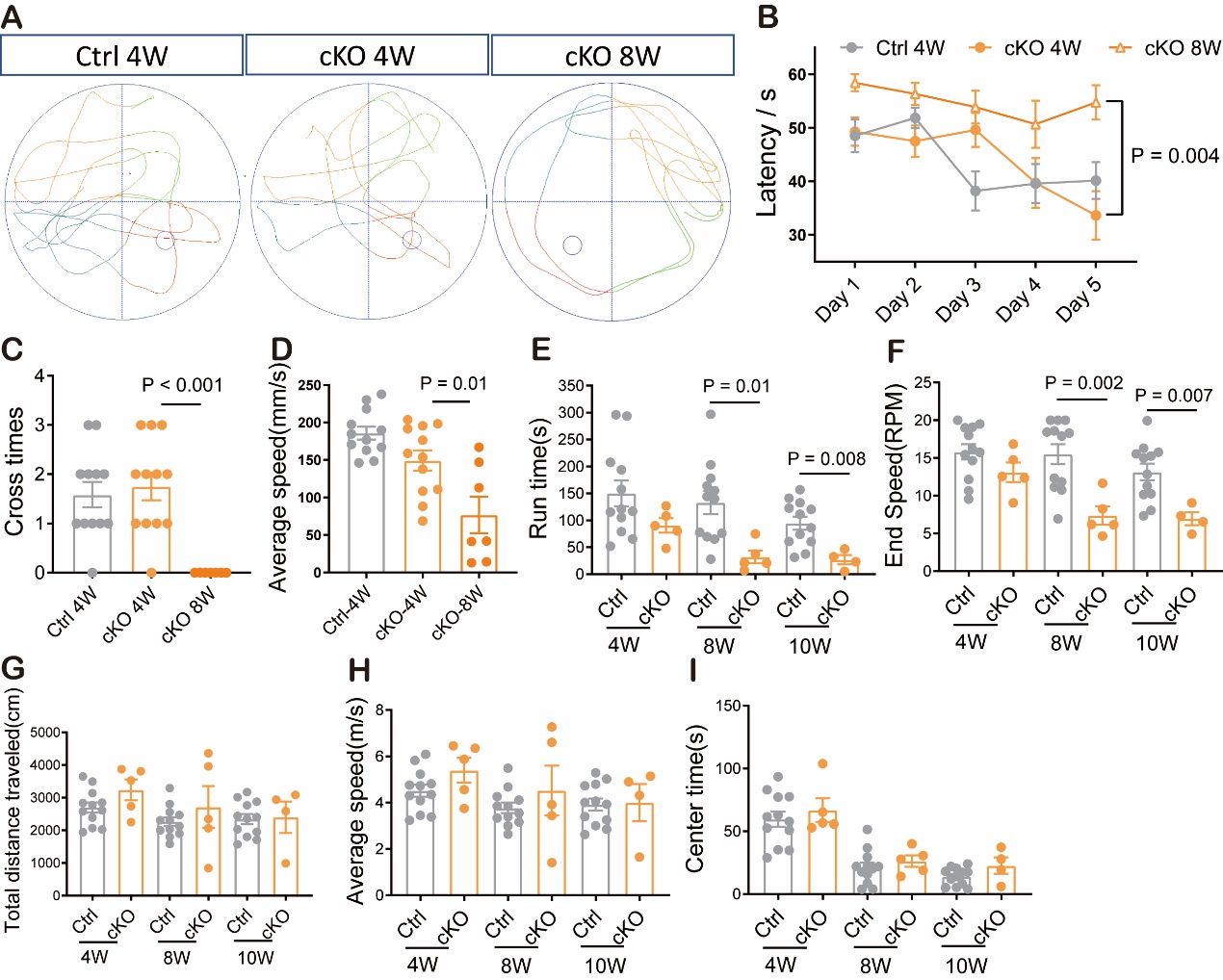

**Figure S3. *Tpk* cKO mice exhibit cognitive and motor dysfunctions.** A. Representative plots depicting the paths of the control (Ctrl) and cKO mice swimming to find the platform in the Morris water maze test at the 4^th^ (4W) and 8^th^ (8W) weeks after tamoxifen treatment. B. Quantification of latency to mount the platform in the Morris water maze across the 5 days training period of the control littermates (4W, n = 12) and cKO mice (4W, n = 11; 8W, n = 7). There were no significant differences at the 4^th^ week between the genotypes. Significant increases were detected in the cKO mice at the 8^th^ week as compared to the 4^th^ week after tamoxifen treatment (P = 0.004). C & D. Quantifications on times of crossing the target site (C) and average velocity in the water maze (D) after retrieval of the platform on testing day. There were no significant differences at the 4^th^ week between the two genotypes, but significant decreases in the cKO mice at the 8^th^ week compared to the 4^th^ week after tamoxifen treatment. E & F. Results on run times (E) and end speed (F) of rotarod test. There were no significant changes in the cKO mice (n = 5) compared to the control littermates (n = 12 for all weeks) at the 4^th^ week (P > 0.05). Significant decreases were detected at the 8^th^ and 10^th^ weeks after tamoxifen treatment. G-I. Results in total distance (Panel G), average speed (Panel H), and center times (Panel I) of open field test. No significant changes were found in the cKO mice (n = 5) at the 4^th^, 8^th^, and 10^th^ weeks as compared to the control littermates (n = 12, P > 0.05).

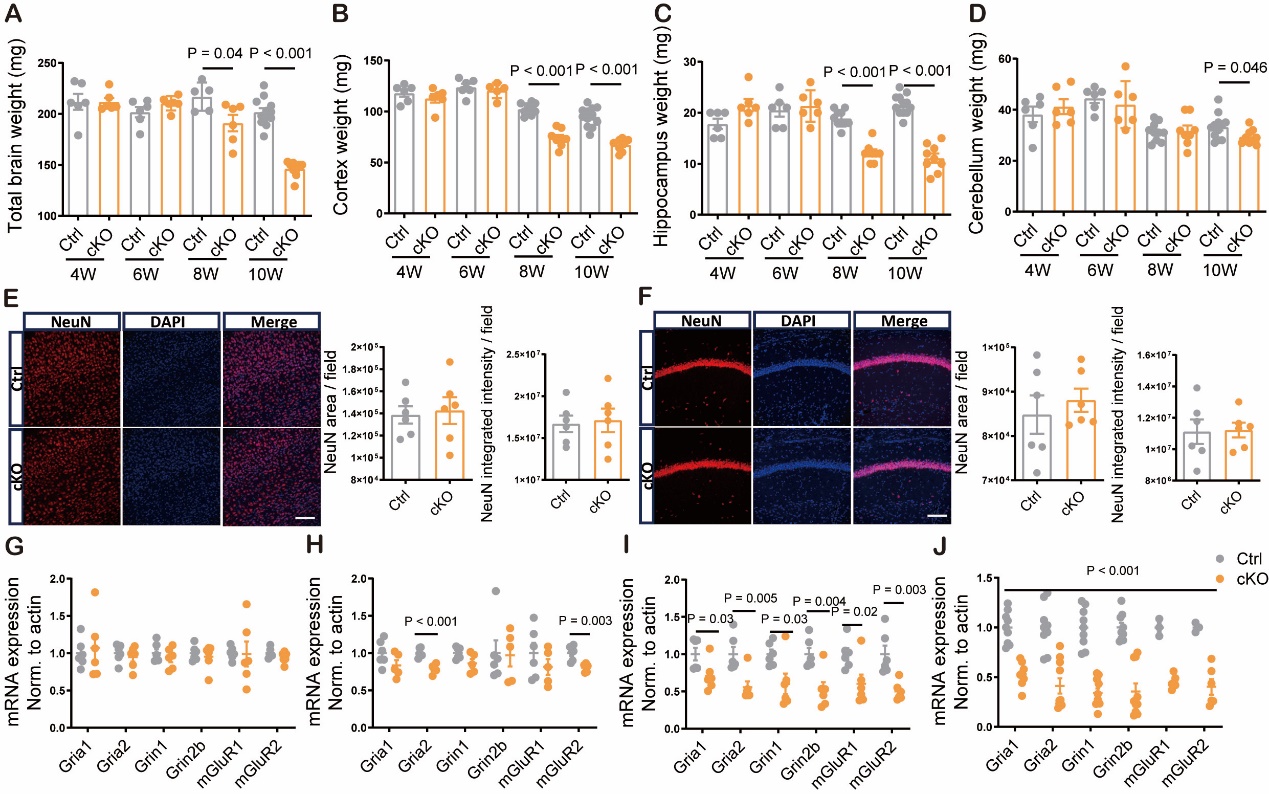

**Figure S4. *Tpk* cKO mice display progressive losses of synapses and neurons as well as brain atrophy.** A-D. Quantifications for the weights of total brain (A), cerebral cortex (B), hippocampus (C), and cerebellum (D) of the control littermates and cKO mice at the 4^th^ (n = 6, 5 for Ctrl and cKO, and the same convention herein), 6^th^ (n = 5, 6), 8^th^ (n = 12, 7), and 10^th^ (n = 9, 7) weeks after tamoxifen treatment. There were no significant differences at the 4^th^ and 6^th^ weeks between the two genotypes (P > 0.05 for all). There were significant decreases in weights of total brain, cerebral cortex, and hippocampus in the cKO mice compared to the control littermates at the 8^th^ and 10^th^ weeks. The weight of cerebellum in the cKO mice was slightly decreased as compared with that in the control littermates only at the 10^th^ week (P = 0.046). E & F. There were no significant differences in cortical (E) and hippocampal (F) NeuN-positive cells between the control littermates (n = 6) and cKO mice (n = 6, P > 0.05) at the 4^th^ week after tamoxifen treatment. Representative images (left) and quantifications for the area (middle) and integrated intensity (right). Scare bar, 100µm. G-J. Comparisons of mRNA levels of genes in synaptic glutamate receptor *Gria*1, *Gria*2, *Grin*1, *Grin*2b, *mGluR*1, and *mGluR*2 in cortical samples between the control littermates and cKO mice at the 4^th^ (G, n = 6, 6), 6^th^ (H, n = 6, 5), 8^th^ (I, n = 6, 6), and 10^th^ (J, n = 9, 10) weeks after tamoxifen treatment. Results of mRNA levels detected by RT-qPCR were normalized to that of β-actin. There were no significant changes in mRNA levels of all the genes tested at the 4^th^ week between the cKO mice and the control littermates (P > 0.05). The mRNA levels of *Gria*2 (P < 0.001) and *mGluR*2 (P = 0.003) in the cKO mice were significantly decreased but other genes were not as compared with that in the control littermates at the 6^th^ week. The mRNA levels of all the genes tested in the cKO mice were significantly decreased as compared with that in the control littermates at the 8^th^ (P < 0.05) and 10^th^ (P < 0.01) weeks.

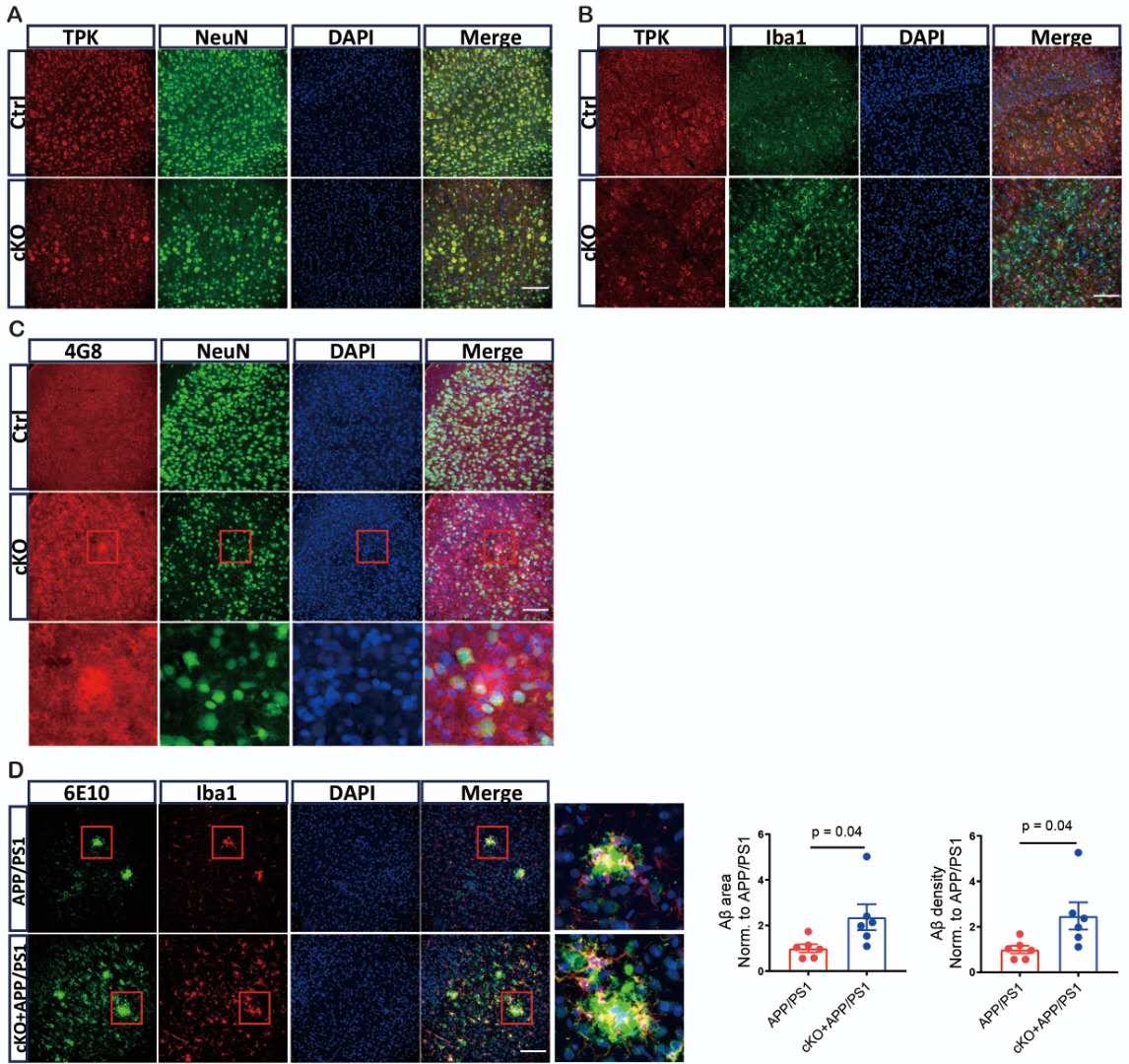

**Figure S5. TPK location and Aβ plaques in mouse brains with conditional *Tpk* knockout.** A. Representative images of TPK and neuron immunohistochemical staining. The results showed that TPK mainly co-existed with neurons stained by NeuN antibody in the control littermates. B. Representative images of TPK and microglia immunohistochemical staining. The results showed that TPK did not co-exist with microglia stained by Iba1 antibody. C. Representative images of Aβ and neurons immunohistochemical staining. The results showed that no Aβ plaques were detected using 4G8 antibody and Aβ did not co-exist with neurons stained by NeuN antibody in brain samples of the cKO mice at 10^th^ weeks after tamoxifen treatment. D. The effect of TPK deficiency on Aβ plaques. Representative images (left) and quantification of Aβ plaques (right) were shown. Aβ plaques represented by Aβ area and density were significantly increased in the cortex of the cKO/APP/PS1 mice as compared to the APP/PS1 and surrounded with microglia at the 6^th^ week after tamoxifen treatment (n = 6, 6, P = 0.04). Scare bar, 100µm.

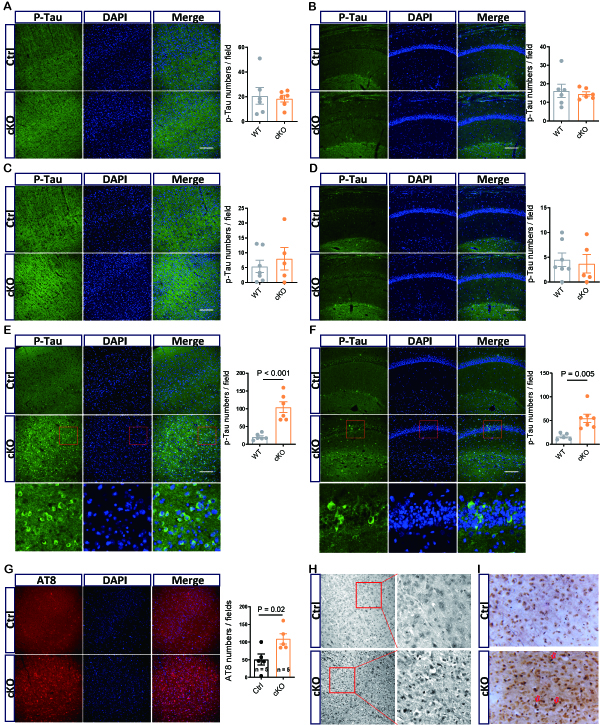

**Figure S6. Tau phosphorylation and tangles in brains of *Tpk* cKO mice.** A-F. The levels of Tau phosphorylation in the cortex (A, C & E) and hippocampum (B, D & F) of control littermates and cKO mice at the 4^th^ (A & B), 6^th^ (C & D), and 8^th^ (E & F) weeks after tamoxifen treatment. Representative images (left) and quantifications of phosphorylated Tau (right) are shown. No significant changes were found in either cortex or hippocampus of the cKO mice compared to the control littermates at the 4^th^ and 6^th^ weeks (n = 6, 6, P > 0.05 for both weeks). There were significant increases in Tau phosphorylation detected by immunofluorescent staining using S396 antibody in cortex (P < 0.001) and hippocampus (P = 0.005) of the cKO mice (n = 7, 5) at the 8^th^ weeks after tamoxifen treatment. G. The neurofibrillary tangles assayed by AT8 antibody in the cortex of the cKO mice were significantly enhanced compared to the control littermates at the 10^th^ week after tamoxifen treatment (n = 5, 5, P < 0.05). Representative images (left) and quantifications of phosphorylated Tau (right) are shown. H. Representative images of phosphorylated Tau assessed by AT8 antibody using immunohistochemical staining. I. Representative images of tangles assessed by silver staining. Scare bar, 100µm.

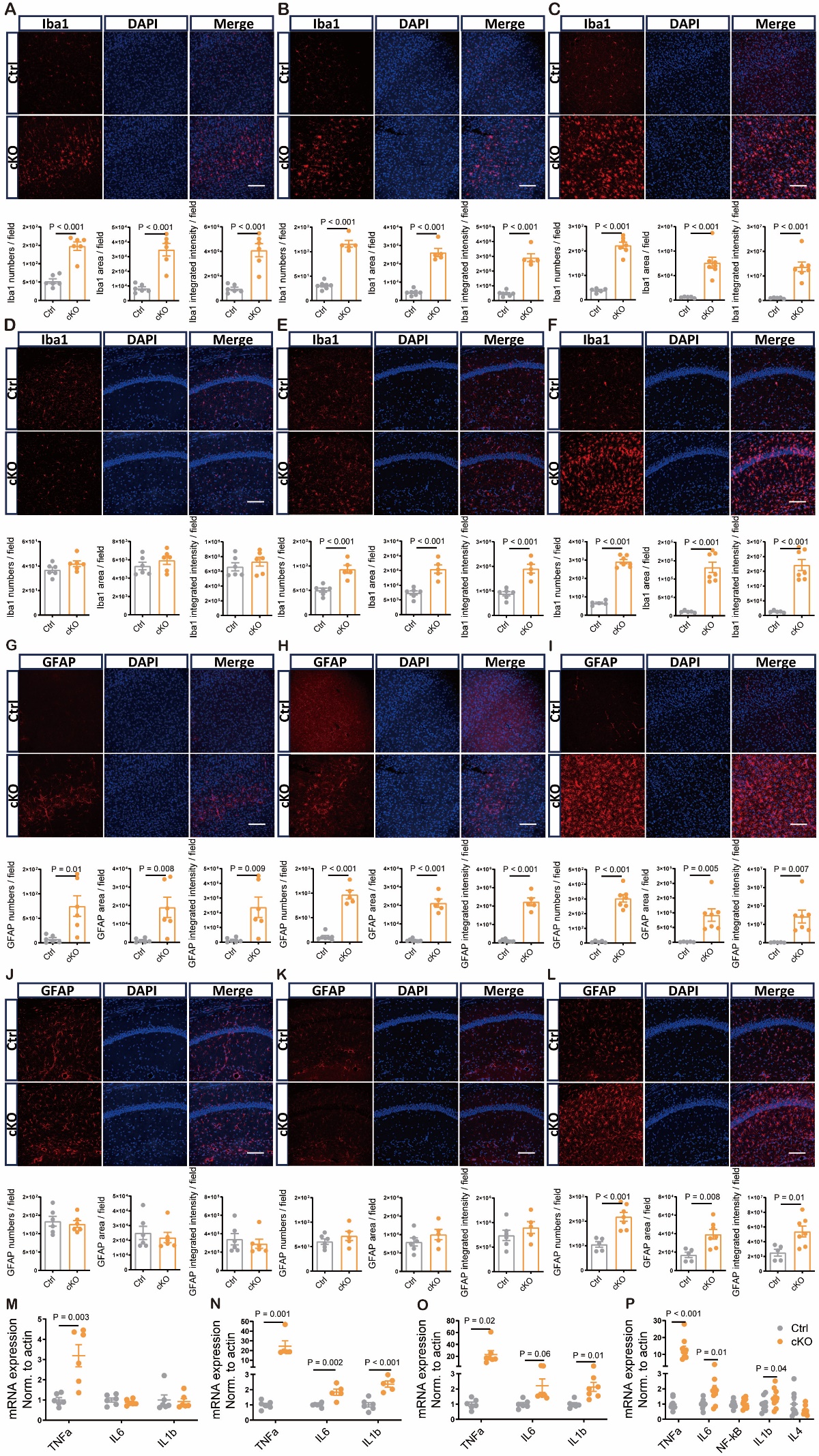

**Figure S7. Glial activation and alteration of inflammatory factors in brains of *Tpk* cKO mice.** A-C. Cortical microglia in the control littermates and cKO mice at the 4^th^ (A, n = 6, 6 for Ctrl and cKO, and the same convention herein), 6^th^ (B, n = 6, 5), and 8^th^ (C, n = 5, 7) weeks after tamoxifen treatment. Representative images (upper panels) and quantifications on the numbers, areas, and intensities of Iba1-positive cells (lower panels) were shown. Significant cortical microglial activation in the cKO mice began at the 4^th^ week and maintained throughout the entire study period as compared with that in the control littermates (P < 0.001 for all). D-F. Hippocampal microglia in the control littermate and cKO mice at the 4^th^ (D, n = 6, 6), 6^th^ (E, n = 6, 5), and 8^th^ (F, n = 5, 7) weeks after tamoxifen treatment. Significant hippocampal microglial activation in the cKO mice was detected at the 6^th^ and 8^th^ weeks (P < 0.001 for all), but not at the 4^th^ week as compared with that in the control littermates (P > 0.05 for all). G-I. Cortical astrocytes in the control littermates and cKO mice at the 4^th^ (G, n = 6, 6), 6^th^ (H, n = 7, 5), and 8^th^ (I, n = 5, 7) weeks after tamoxifen treatment. Representative images (upper panels) and quantifications on the numbers, areas, and intensities (lower panels) of GFAP-positive cells were shown. Significant cortical astrocyte activation in the cKO mice began at the 4^th^ week and maintained throughout the entire study period (P ≤ 0.01 for all) as compared with that in the control littermates. J-L. Hippocampal astrocytes in the control littermate and cKO mice at the 4^th^ (J, n = 6, 6), 6^th^ (K, n = 6, 5) and 8^th^ (L, n = 5, 7) weeks after tamoxifen treatment. Significant hippocampal astrocyte activation was detected at the 8^th^ week (L, P ≤ 0.01 for all), but not at the 4^th^ and 6^th^ weeks (P > 0.05 for all) in the cKO mice compared to the control littermates. Scare bar, 100µm. M-P. The mRNA levels of inflammatory cytokines measured by RT-qPCR in brain samples of the control littermates and cKO mice at the 4^th^ (M, n = 6, 6), 6^th^ (N, n = 6, 5), 8^th^ (O, n = 6, 7), and 10^th^ (P, n = 9, 10) weeks after tamoxifen treatment. The mRNA levels of TNFα were significantly increased in the cKO mice at all test points (P < 0.005 for all). The mRNA levels of IL-6 were significantly increased in the cKO mice at the 6^th^ and 10^th^ weeks (P < 0.05 for both), but not at the 4^th^ and 8^th^ weeks (P > 0.05 for both) as compared with that in the control littermates. The mRNA levels of IL-1b were significantly increased in the cKO mice at the 6^th^, 8^th^, and 10^th^ weeks (P < 0.05 for all), but not at the 4^th^ week (P > 0.05) as compared with that in the control littermates. No significant changes were detected in the mRNA levels of NF-κB and IL-4 (P > 0.05) in the cKO mice even at the 10^th^ week after tamoxifen treatment as compared with that in the control littermates (P).

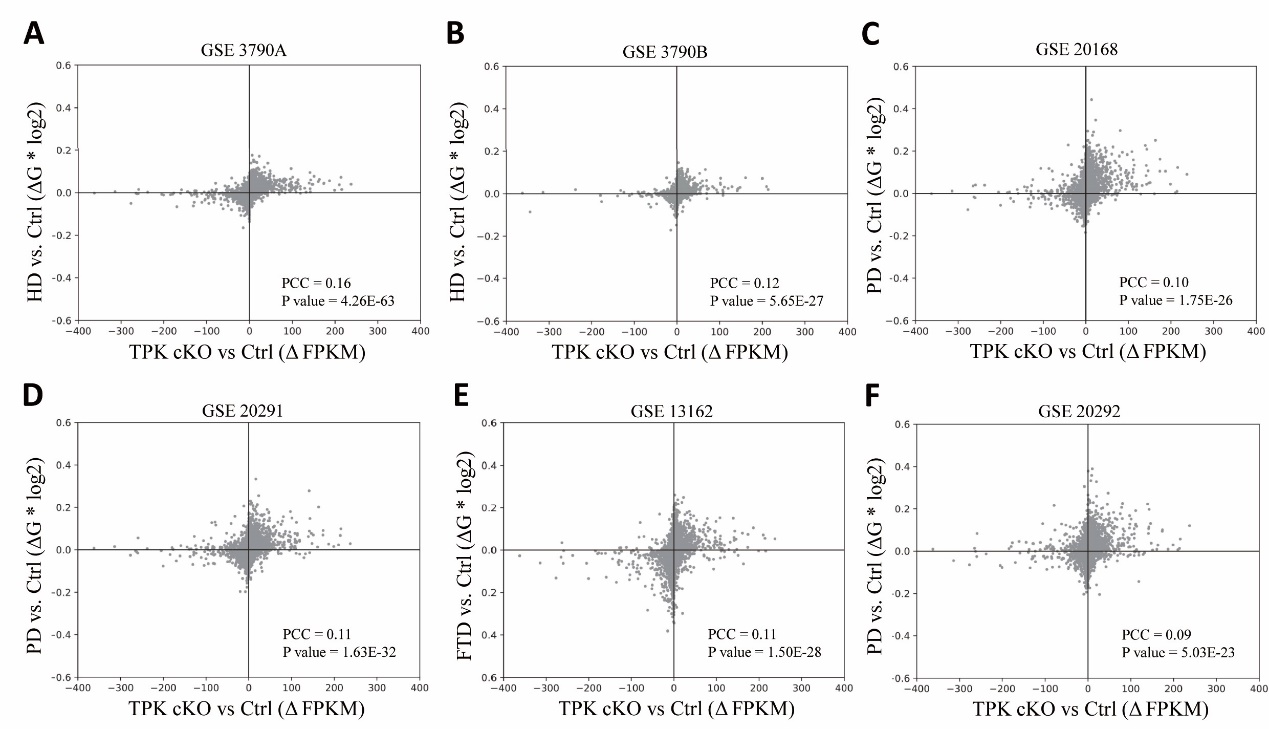

**Figure S8. Comparisons of cross species analysis of expression changes in *Tpk* cKO mice with those in patients with Huntingdon’s disease, Parkinson’s disease, and frontotemporal lobar degeneration.** Scatter plots showing genome-wide correlated changes in gene expression between the cortical samples of the cKO mice and brain samples of patients with Huntingdon’ disease (HD), Parkinson’s disease (PD), and frontotemporal lobar degeneration (FTLD). A: The cKO mice and HD-GSE3790A (Frontal cortex, cerebellum, and caudate nucleus), PCC = 0.16, P = 4.23e-63; B: The cKO mice and HD-GSE3790B (Frontal cortex, cerebellum, and caudate nucleus), PCC = 0.12, P = 5.65e-27; C: The cKO mice and PD-GSE20168 (Prefrontal cortex), PCC = 0.10, P = 1.75e-26; D: The cKO mice and PD-GSE20291 (Putamen nucleus): ACC = 0.11, P = 1.63e-32; E: The cKO mice and FTLD-GSE13162 (Frontal cortex): ACC = 0.11, P = 1.5e-28; F: The cKO mice and PD-GSE20292 (Substantia nigra): ACC = 0.09, P = 5.03e-23.

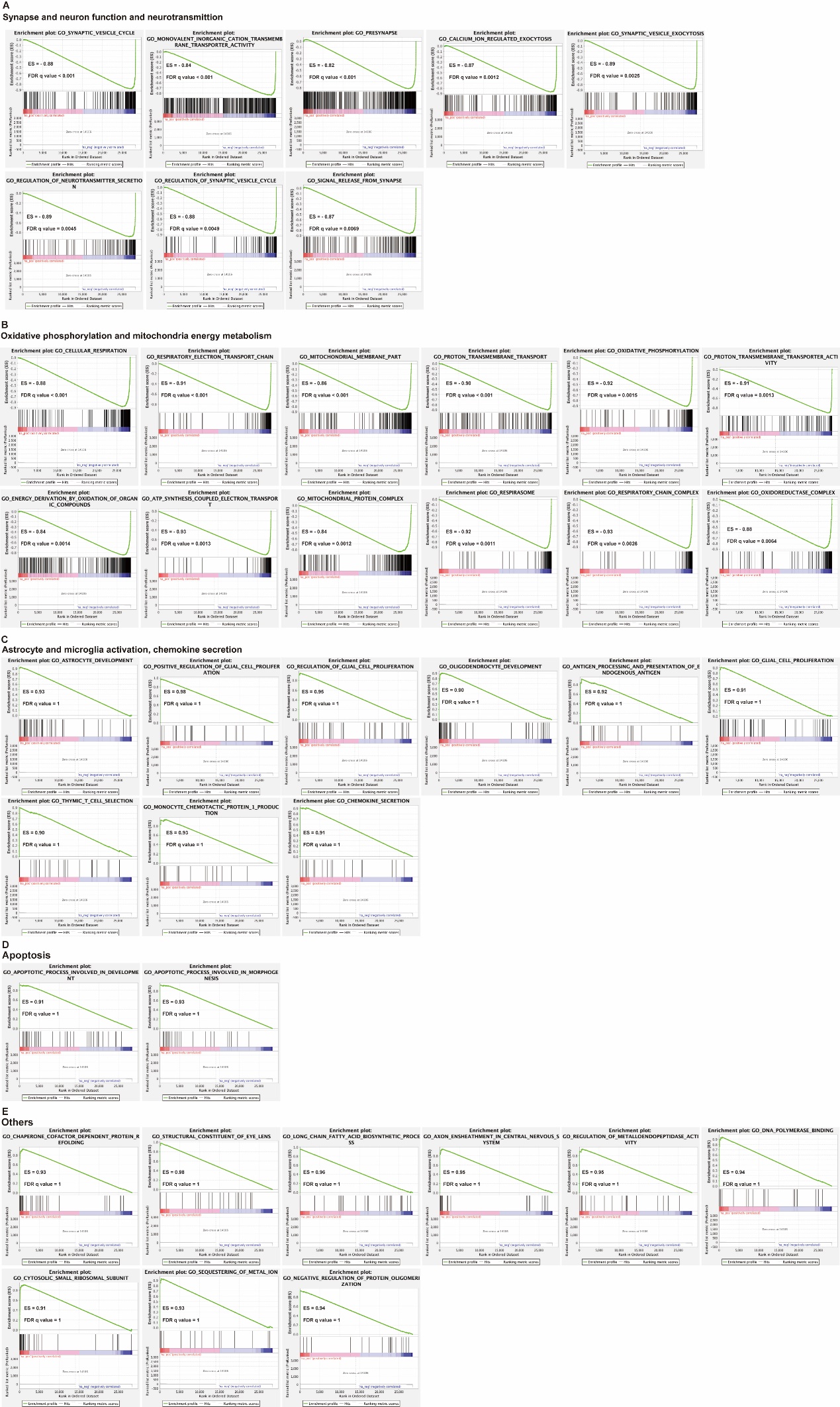

**Figure S9. Enriched pathways in RNAseq data of GSE95587 dataset for AD fusiform gyrus samples.** Gene set enrichment plots of pathways where AD impacted transcripts (x-axis) are sorted by magnitude of upregulation (red) to downregulation (green). A. Gene sets showed significant down-regulation of synaptic, neuronal, and neurotransmittal functions, including synaptic vesicle cycle, monovalent inorganic cation transmembrane transporter activity, presynapse, calcium on regulated exocytosis, synaptic vesicle exocytosis, regulation of neurotransmitter secretion, regulation of synaptic vesicle cycle, and signal release from synapse. B. Gene sets showed significant down-regulation of oxidative phosphorylation and mitochondria energy metabolism, including cellular respiration, respiratory electron transport chain, mitochondrial membrane part, proton transmembrane transport, oxidative phosphorylation, proton transmembrane transporter activity, energy derivation by oxidation of organic compounds, ATP synthesis coupled electron transport, mitochondrial protein complex, respirasome, respiratory chain complex, and oxidoreductase complex. C. Gene sets showed significant up-regulation of astrocyte/microglia activation and chemokine secretion, including astrocyte development, positive regulation of glial cell proliferation, regulation of glial cell proliferation, oligodendrocyte development, antigen processing and presentation of endogenous antigen, glial cell proliferation, thymic T cell selection, and monocyte chemotactic protein 1 production, chemokine secretion. D. Gene sets showed significant up-regulation of apoptosis, including apoptotic process involved in development and morphogenesis. E. Gene sets showed significant up-regulation of others, including chaperone cofactor dependent protein refolding, structural constituent of eye lens, long chain fatty acid biosynthetic process, axon ensheathment in central nervous system, regulation of metalloendopeptidase activity, DNA polymerase binding, cytosolic small ribosomal subunit, sequestering of metal ion, and negative regulation of protein oligomerization.

**Table S1**. **Specially down-regulated TPK in Alzheimer’s disease**

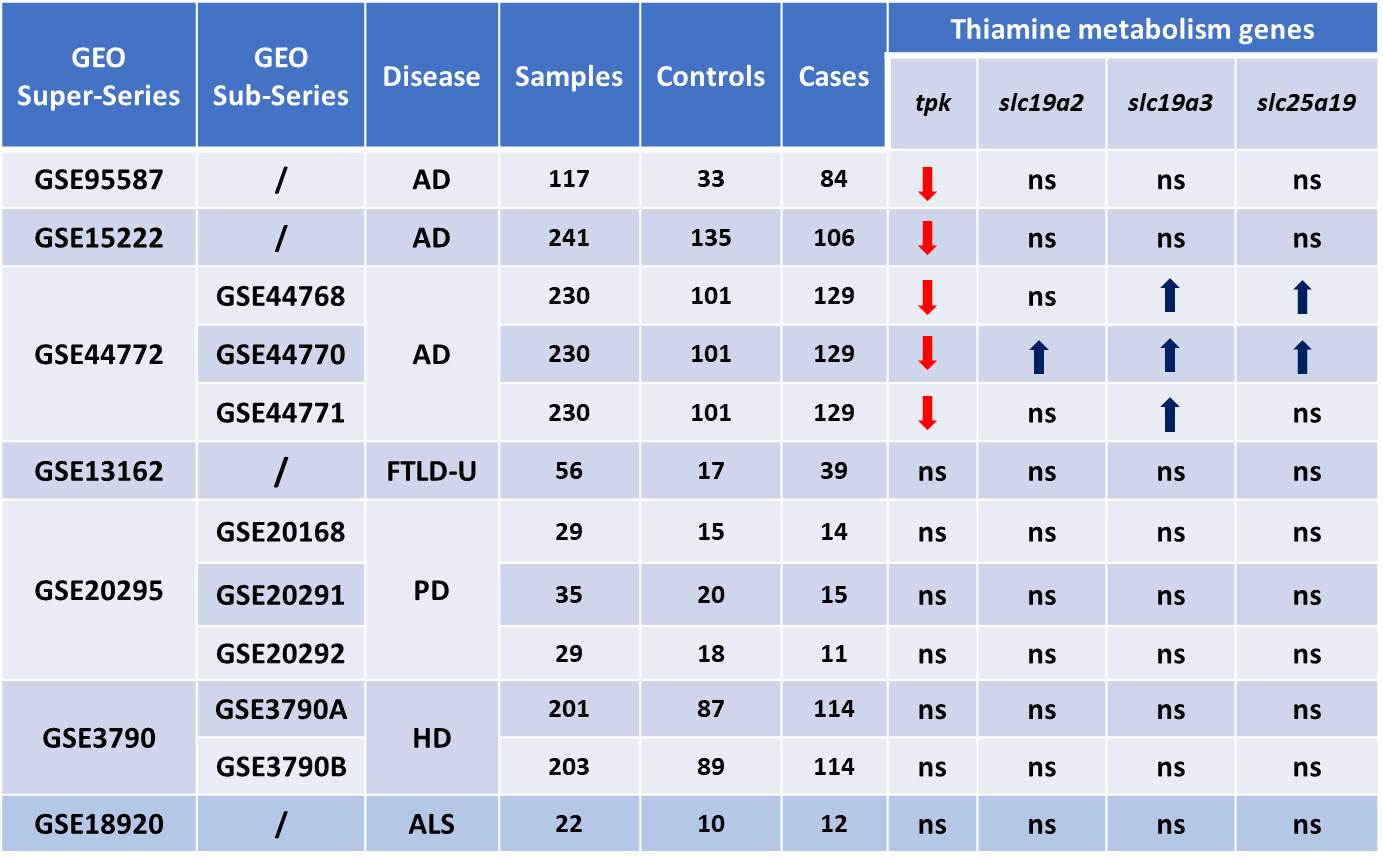

(**↓**indicates significant reduction with P < 0.01, **↑**indicates significant increase with P < 0.05, ns indicates no significant change)

**Table S2**. **The results of cross-species correlation analysis in genome-wide transcription levels between the cKO mice and AD patients.**

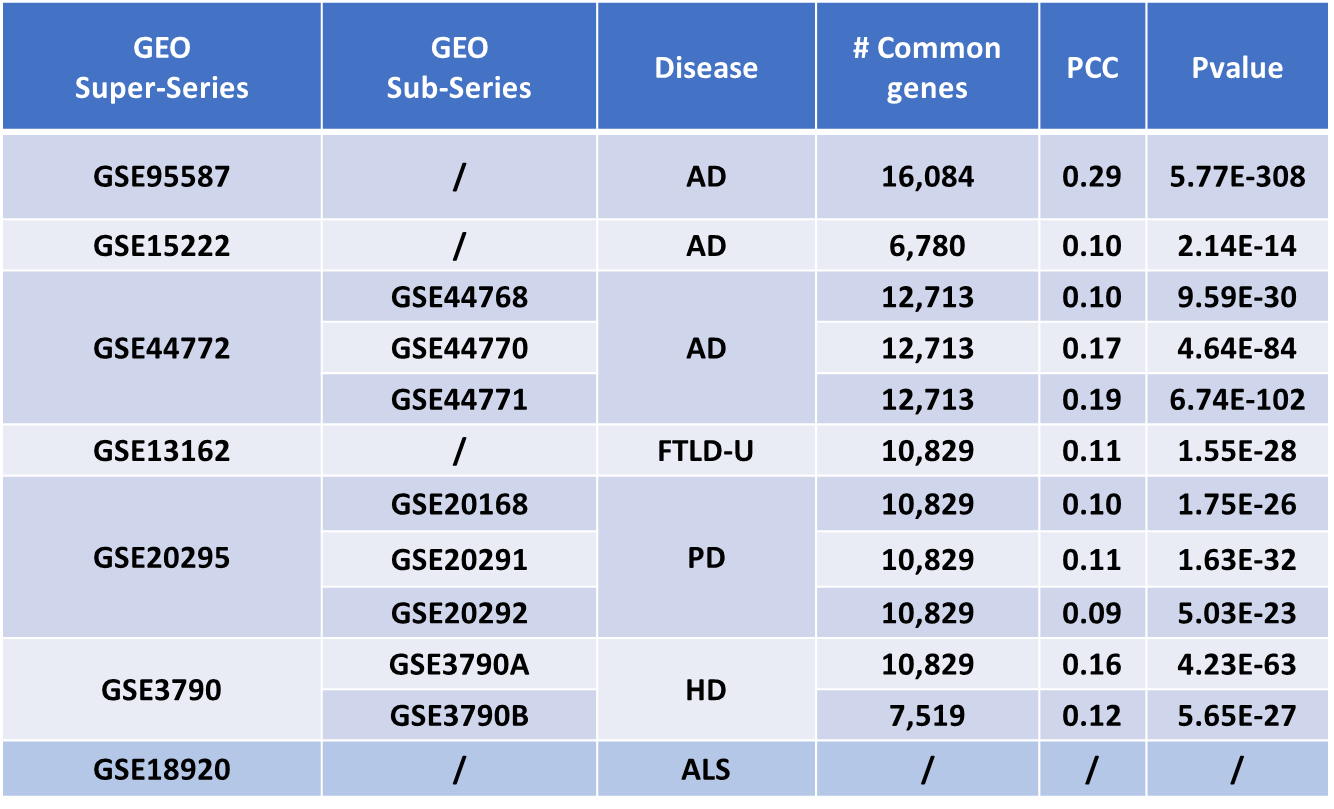
