## Supplementary Materials and methods for "Thiamine pyrophosphokinase deficiency induces Alzheimer’s pathology"

**Methods and Materials**

**GEO datasets download**

To detect the expression changes of all known genes associated thiamine metabolism, including *thiamine pyrophosphokinase 1* (*TPK*), *SLC19A2*, *SLC19A3*, and *SLC25A19*, a total of 7 public RNA-seq and microarray datasets for brain samples of patients with Alzheimer’s disease (AD), Parkinson’s disease (PD), frontotemporal lobar degeneration (FTLD), Huntington’s disease (HD), and amyotrophic lateral sclerosis (ALS) from the GEO databases were analyzed. The datasets are GSE13162, GSE20295, GSE95587, GSE15222, GSE18920, GSE3790, and GSE44772.

GSE95587 is an Illumina HiSeq 2500 RNA-seq dataset, measuring fusiform gyrus tissue samples of 117 subjects, including 84 AD patients and 33 neurologically normal age-matched control subjects This dataset was downloaded from <https://www.ncbi.nlm.nih.gov/geo/download/?acc=GSE95587&format=file>.

GSE44772 is a Rosetta/Merck Human 44k 1.1 microarray dataset, which is composed of the following Sub-Series: GSE44768, GSE44770 and GSE44771.

GSE44768 measures cerebellar tissue samples of 230 subjects, including 129 late onset AD (LOAD) patients and 101 non-demented healthy controls. This dataset was downloaded from ftp://ftp.ncbi.nlm.nih.gov/geo/series/GSE44nnn/GSE44768/ matrix.

GSE44770 measures tissue samples of dorsolateral prefrontal cortex in the same 230 subjects. This dataset was downloaded from ftp://ftp.ncbi.nlm.nih.gov/geo/series/ GSE44nnn/GSE44770/matrix.

GSE44771 measures tissue samples of visual cortex in the same 230 subjects. This dataset was downloaded from ftp://ftp.ncbi.nlm.nih.gov/geo/series/GSE44nnn/ GSE44771/matrix.

GSE15222 is a Sentrix Human Expression BeadChip dataset, measuring postmortem brain samples of 363 subjects, including 176 late-onset AD (LOAD) cases and 187 neuropathologically normal controls. Due to the datasets were mixed with data from multiple brain regions, only 135 neuropathologically normal controls and 106 AD cases temporal cortex datasets were used for further analysis (Table).

This dataset was downloaded from [ftp://ftp.ncbi.nlm.nih.gov/geo/series/GSE15nnn/ GSE15222/matrix](ftp://ftp.ncbi.nlm.nih.gov/geo/series/GSE15nnn/%20GSE15222/matrix).

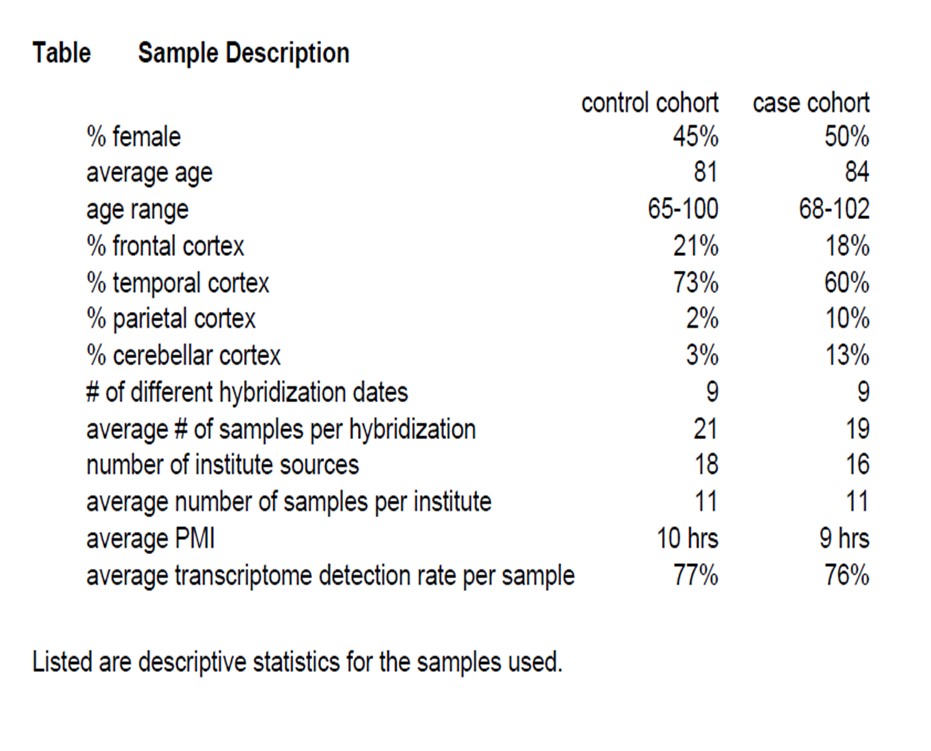

GSE13162 is an Affymetrix Human Genome U133A 2.0 Array dataset, measuring postmortem frontal samples of 56 subjects, including 39 FTLD patients and 17 normal controls. This dataset was downloaded from https://www.ncbi.nlm.nih.gov/geo/ download/?acc=GSE13162&format=file.

GSE20295 is an Affymetrix Human Genome U133A Array dataset, which is composed of the following Sub-Series: GSE20168, GSE20291 and GSE20292.

GSE20168 measures prefrontal tissue samples of 29 subjects, including 14 PD patients and 15 normal controls. This dataset was downloaded from <https://www.ncbi.nlm.nih.gov/geo/download/?acc=GSE20168&format=file>.

GSE20291 measures putamen tissue samples of 35 subjects, including 15 PD patients and 20 normal controls. This dataset was downloaded from https://www.ncbi. nlm.nih.gov/geo/download/?acc=GSE20291&format=file.

GSE20292 measures substantia nigra tissue samples of 29 subjects, including 18 PD patients and 11 normal controls. This dataset was downloaded from https://www.ncbi. nlm.nih.gov/geo/ download/?acc=GSE20292&format=file.

GSE3790 is an Affymetrix Human Genome Array dataset. This dataset was downloaded from https://www.ncbi.nlm.nih.gov/geo/download/?acc=GSE3790& format=file. Samples of this dataset were profiled using two different Array platforms (Affymetrix Human Genome U133A Array and Affymetrix Human Genome U133B Array). Therefore, samples were separately analyzed in different platforms. The dataset from Affymetrix Human Genome U133A Array was named GSE3790A, and the dataset from Affymetrix Human Genome U133B Array was named GSE3790B. GSE3790A measures postmortem frontal cortex, cerebellum, and caudate nucleus samples of 201 subjects, including 114 HD patients and 87 normal controls. GSE3790B measures postmortem frontal cortex, cerebellum, and caudate nucleus samples of 203 subjects, including 114 HD patients and 89 normal control subjects.

GSE18920 measures 22 total lumbar spinal cords samples including 12 sporadic ALS and 10 control subjects. Because no annotation files could be identified for this dataset, we obtained expression data for *TPK*, *SLC19A2*, *SLC19A3*, and *SLC25A19* from the well analyzed datasets (http://research-pub.gene.com/BrainMyeloidLandscape/) ^26^. However, cross-species correlation analysis could not be performed for this dataset.

**GEO data processing**

Affymetrix microarray data GSE13162, GSE20168, GSE20291, GSE20292, GSE3790A, and GSE3790B were processed (including background correction, normalization, and expression quantification) using the Robust Multi-Array Average (RMA) method implemented in R package *affy*1 (version 1.48.0). For GSE13162, GSE20168, GSE20291, GSE20292, and GSE3790A, gene annotation files were downloaded from <https://www.ncbi.nlm.nih.gov/geo/query/acc.cgi?acc=GPL96>, and for GSE3790B, the gene annotation file was downloaded from https://www.ncbi.nlm.nih.gov/geo/query/acc.cgi?acc=GPL97. Expression levels of multiple probe sets corresponding to the same gene were averaged to represent the expression level of the gene.

**Illumina HiSeq 2500 data processing**

Data of GSE95587 have already been processed using "nRPKM" normalization. This is similar to RPKM normalization but includes an extra step for size factor adjustment as previously described.

**Sentrix Human Expression BeadChip data processing**

We used the processed data with probe set expression levels of GEO15222. Missing values were imputed with the mean value across all samples using locally developed Perl code. The gene annotation file was downloaded from https://www.ncbi.nlm.nih.gov/geo/query/acc.cgi?acc=GPL2700. Expression levels of multiple probe sets corresponding to the same gene were averaged to represent the expression level of the gene. Genes with negative expression values were eliminated.

**Rosetta/Merck Human 44k 1.1 microarray data processing**

Gene expression profiling was provided for GEO44768, GEO44779, and GEO44771. Gene expression was reported as the mean-log ratio of individual microarray intensities relative to average intensities of all samples. Detailed data processing steps were described in the Experimental Procedures of the reference ^27^.

**Cross-species correlation analysis**

All genes in the RNA-seq data of the cKO mice were mapped to human genes using the NCBI homologene database ^28^. Expression levels of mapped genes were then extracted from the origin RNA-seq data. Common genes found between RNA-seq data of the cKO mice and each of the GEO datasets were used to perform cross-species correlation analysis as previously described ^29^. Correlation of the changes in gene expression between the cKO mice and patients of the individual GEO dataset was assessed using Pearson’s correlation coefficient (PCC). For each pair of comparison, a P value was derived from the PCC using Fisher's Z transformation test.

**Animals**

All animal care and experimental procedures were approved by the Medical Experimental Animal Administrative Committee of Fudan University and by the Institutional Animals Care and Use Committee of the Institute of Neuroscience, Shanghai Institutes for Biological Science, Chinese Academy of Science.

**Generation of conditional TPK knockout mice.**

To generate the conditional knockout of mTPK allele (NM_013861.3), for which nine exons have been identified with the ATG start codon in exon 2 and TAA stop codon in exon 9, exon 4 was selected for targeting to generate a conditional knockout mTPK allele (Fig. S2A). The deletion of exon 4 resulted in the loss of function of TPK gene by a frameshift of downstream exons. The detailed procedures are as follows. Vector construction: Mouse genomic fragments were amplified from the BAC clone with high fidelity Taq DNA polymerase and assembled into a targeting vector together with recombination sites and selection markers, as indicated on the vector map, in which the Neo cassette is flanked by Frt sites, CKO region flanked by LoxP sites, and DTA used for negative selection (Fig. S2B). The targeting construct was electroporated into C57BL/6 ES cells. Eighty-five (one 96-well plate) G418-resistant colonies were picked and screened by PCR using the strategy shown (Fig. S2C). The PCR primers were: 3’ arm - F: (GCTGACCGCTTCCTCGTGCTTTA) / 3’ arm - R: (GACCACCAAACAGCATACACCAGCT) and LOXP-F: (GTCACTTGAATATACCACAGTTCCCAG) / LOXP - R: (CATATTGGCACACTA CCCTGGAT). Two potential targeted clones (C10 and E7) were identified (Fig. S2D) and both were further analyzed by Southern blotting (Fig. S2E). The genomic DNA of the potential clones C10 and E7 were digested by Nde I and analyzed by southern blot for a 7.3 kb band from wild type allele and a 9.3 kb band from recombinant allele using probes generated by PCR with primers: 5’- probe, 5’ arm - F: ATCTTTCCAGTGTCTCATTC / 5’ arm - R: TACTTCTTGTTTCGGTCTT; and digested by Xba I and analyzed by southern blot for a 11.1 kb band from wild type allele and a 8.4 kb band from recombinant allele, using probes generated by PCR with primers: 3’- probe, 3’ arm - F: GGATGTTGGCATTGAAGC / 3’ arm - R：GATTTTAGGAGACGAGCA, E7 were positive. The ES cells with the targeting construct confirmed by PCR and Southern analysis were injected into C57BL/6N blastocysts and subsequently transplanted into the uterus of pseudo-pregnant females. Chimeric males generated from albino-B6 blastocysts injected with the mutant ES cells were selected based on coat color and subsequently crossbred with C57BL/6N females. Germline mutant mice were crossbred with ROSA26-FLPe knock-in mice (Jackson Laboratory, stock no. 009086) to remove the FRT-flanked splice acceptor site, and neomycin resistance cassette. TPK flox /+ mice were identified using primers: LoxP_F: GAATTCCAAAGCCAGCATGT / LoxP_R: CCACTCACAGACCCTTTGCA and Neo F: ACTGTGCTTTGCTATCT CTACTGACA / Neo R: GGGCCACAAACACCTTCACATCT. The expected 192 bp fragment from wildtype allele and 253 bp fragment from recombinant allele were represented as positive F1 mice. Four positive pups (2, 4, 5, 6) from clone E7 were identified (Fig. S2F). Resultant TPK flox /+ mice were interbred to generate homozygous mutant mice with conditional knockout potential (TPK flox/flox). The TPK flox/flox mice were crossed with excitatory neuron-specific Cre transgenic mice, CaMKIICre^Ert2/+^ strain (EMMA ID: 02125). To generate brain excitatory neuron-specific TPK knockout, mice at 12 weeks of age were intraperitoneally injected with tamoxifen (50 mg/kg per day) for five consecutive days.

**Immunohistochemical staining**

After deeply anaesthetized with 0.14 g/kg sodium pentobarbital, the mice were intracardially perfused with phosphate buffered saline (PBS) and then 4% paraformaldehyde for fixation. Serial coronal sections (30 μm) were cut with a sliding microtome (Leica) and stained using a freely floating method. For examination of amyloid plaques, the sections were incubated in 88% formic acid at the room temperature (22-25 °C) for 8 min. After being washed in PBS at pH 7.4, the sections were put into PBS containing 5% bovine serum albumin (BSA) and 0.5% Triton X-100 for 2 h at 37 °C in order to block non-specific reactions. Then, the sections were incubated with primary antibodies for NeuN (Millipore, MAB377, 1: 500), GFAP (CST, 3670S, 1: 500), Iba1 (Wako, 019-19741, 1:500), 6E10 (Covance, SIG-39300, 1:1000), 4G8 (Covance, SIG-39220, 1:150) or P-tau (Santa cruz, sc-101815, 1:50) overnight at 4 °C. After being washed with PBS, the sections were incubated with goat anti-mouse antibody conjugated to Alexa Fluor 488 (Invitrogen, 1:500) or Alexa Fluor 546 (1:500, Invitrogen) for 2 h at 37 °C. Nuclei were stained with DAPI (Sigma, D9542, 1:1000) and the slices mounted on 3-aminopropyltriethoxysilane (APES)-coated glass slides. Z-stack images were taken using Nikon A1 (Tokyo, Japan) laser scanning microscope, with a 25 x objective. Results were quantified using ImageProPlus (Media Cybernetics, Silver Spring, MD, USA).

**Protein purification and western blotting**

Brain tissues were lysed with the radioimmunoprecipitation assay (RIPA) lysis buffer containing the protease inhibitor mixture and PhosSTOP phosphatase inhibitor cocktail (Roche). Protein concentrations were determined with the Pierce™ BCA protein assay kit according to the manufacturer’s instruction (Thermo Scientific, Rockford, IL). The same amount of proteins was loaded in each lane for electrophoresis on a 10% denaturing Tris/glycine sodium dodecyl sulfate (SDS) polyacrylamide gel and subject to electrophoresis. Proteins were transferred to the polyvinylidene fluoride membranes (Millipore) and blocked in 5% milk in Tris-buffered saline supplemented with 0.1% Tween-20 (TBS-T, pH 7.4) for 2 h. The membrane incubated overnight at 4^o^C with TBST containing 3% milk and primary antibodies for TPK1 (Proteintech, 10942-1-AP, 1:1,000), beta-Actin (Proteintech, 60008-1-Ig, 1:10,000), APP (BOSTER, PB0101, 1:1,000), or BACE1 (CST, 5606, 1:1000). Then, membranes were washed in TBS-T and incubated for 2 h with horseradish peroxidase (HRP)-conjugated anti-mouse or anti-rabbit IgG (Millipore) in TBS-T containing 3% milk. After a final wash in TBST, member were incubate with ECL substrate (Pierce® Fast Western Blot Kit, 35050, USA) for 1–2 minutes, signals were detected with Tanon 6200 Luminescent Imaging Workstation (Tanon Science & Technology Co., Shanghai, China). Results were quantified using Image J (N.I.H.).

Frozen human brain tissue homogenates in RIPA buffer (1% Triton X100, 1% sodium deoxycholate, 0.2% SDS, 0.15M NaCl, 0.05M Tris-HCl pH 7.2) supplemented with complete mini protease inhibitor cocktail tablet (Roche, 11836170001) were resolved by SDS-PAGE on 10% tris-glycine gels, and then transferred to nitrocellulose membranes. Membranes were blocked in 5% skim milk at room temperature for 1hour, and sequentially blotted with primary antibodies (Abcam, ab249546) and IRDye fluorescence labelled secondary antibodies (LI-COR Biosciences, 926-68021). Actin be used as internal control (Sigma, A5441).

**Real-time qPCR**

Total RNA was extracted from brain tissues of mice using TRIzol reagent (Invitrogen). For Real-time RT-PCR experiments, complimentary cDNA was synthesized using the PrimeScript™ RT reagent Kit with gDNA Eraser (TaKaRa, Japan) according to the manufacturer's protocol. Primer design were using online Primer-BLAST software according to the instructions：[https://www.ncbi.nlm.nih.gov/tools/ primer-blast](https://www.ncbi.nlm.nih.gov/tools/%20primer-blast)/index.cgi?LINK_LOC=BlastHome. Real-time PCR was performed using SYBR Green Master Mix (TaKaRa, Japan) on 7500 Real-time PCR system (Applied biosystems). All reactions were performed in triplicates, and the results normalized to that of control groups from the same preparation.

| Primer | Forward | Reverse |
| --- | --- | --- |
| *β-Actin* | AGAAGGACTCCTATGTGGGTGA | CATGATCTGGGTCATCTTTTCA |
| *Il-1β* | GCAACTGTTCCTGAACTCAACT | ATCTTTTGGGGTCCGTCAACT |
| *Il-4* | GGTCTCAACCCCCAGCTAGT | GCCGATGATCTCTCTCAAGTGAT |
| *Il-6* | GTCCTTCCTACCCCAATTTCCA | TAACGCACTAGGTTTGCCGA |
| *Nf-κb* | ATGGCAGACGATGATCCCTAC | TGTTGACAGTGGTATTTCTGGTG |
| *Tnf-α* | CCCTCACACTCAGATCATCTTCT | GCTACGACGTGGGCTACAG |
| *Gria*1 | CGAGTTCTGCTACAAATCCCG | TGTCCGTATGGCTTCATTGATG |
| *Gria*2 | AAAGAATACCCTGGAGCACAC | CCAAACAATCTCCTGCATTTCC |
| *Grin*1 | CGGCTCTTGGAAGATACAG | GAGTGAAGTGGTCGTTGG |
| *Grin*2b | TTTGGAGATGGGGAGATGG | CAGACACCCATGAAGCAATG |
| *mGluR*1 | AGTCTGCAGAACCGTCTGTG | GTTTACGGGACCTCTCAGGG |
| *mGluR*2 | GACTCTGGCTCCACTAAAGA | AGTCCTCACAAACACAGAGG |

**RNA-seq analysis**

RNA samples extracted from brain cortex of mice as described above were used. Short read FASTQ files were quality trimmed using FASTX toolkit (v. 0.0.14) to trim three bases from the 5′ end of the reads. Paired-end reads were then mapped to the mm9 genome using tophat245 and the UCSC knownGene gtf file. The following parameters were used in the tophat2 call “-N 1 –g 1 –readgap-length 1 –mate-inner-dis 170”. Reads that had the same starting location and strand with mate-pairs that also had the same location and strand were considered to be PCR duplicates and removed from subsequent analyses using Picard tools (v. 1.103). Differentially expressed transcripts were determined using Cufflinks and Cuffdiff (v2.1.1). Downstream analyses were performed in R/Bioconductor47 and used gene summarized expression levels normalized using Fragments Per Kilobase per Million (FPKM) from Cufflinks. Hierarchical clustering was performed using the pvclust R package, where significance was determined using bootstrapping. Principle Components Analysis (PCA) was conducted using the “prcomp” function of the stats package in R/ Bioconductor. Enriched gene ontologies were determined using the package “GOstats” (v. 3.1.1). Gene Set Enrichment Analysis (GSEA) was performed using a pre-ranked gene list determined by cuffdiff and GSEA (v. 2.1.0). Hierarchical clustering of gene expression data was performed using average clustering in the heatmap.2 package. UCSC-style displays of gene expression data were plotted using the “rtracklayer” package and custom R scripts to display RNA sequencing reads as histograms.

**ELISA for Aβ42 and Aβ40**

The levels of Aβ42 and Aβ40 were determined by ELISA (Mouse Aβ42 or Aβ40 Colorimetric ELISA, KMB3441 or KMB3481, Invitrogen, Grand Island, NY, USA) according to the manufacturer’s instructions. In brief, brain tissues were weighed and homogenized in ice-cold PBS containing the protease inhibitor cocktail (Complete Protease Inhibitor Cocktail, Roche Diagnostics) followed by centrifugation at 16,000 x g for 20 min at 4°C, guanidine buffer (5 M guanidine HCl/50 mM Tris-HCl, pH 8.0). The homogenates were mixed for 4 hours at the room temperature and were diluted 1:5 in PBS containing 5% BSA and 0.03% Tween-20 supplemented with the protease inhibitor cocktail followed by centrifugation at 16,000 x g for 20 min at 4°C. The supernatant was diluted and analyzed according to the manufacturer’s instructions. Final values of Aβ are expressed as pg per gram of brain tissue (wet weight).

**Y-maze test**

The Y-maze consists of 3 black horizontal arms (36 cm long, 5 cm wide, and 10 cm high) at 120° angles to each other. Mice were allowed to move freely through the Y-maze during a 5-min session. Alternation was defined as successive entries into the three arms on overlapping triplet sets. The percentage of alternation was calculated as the total number of alternations ×100 / (total number of arm entries - 2), which is not influenced by the unwillingness of the mouse to move. The performance was video-recorded and analyzed by an experimenter who was blinded with the genotypes of the animals using the image analyzing software (ANY-maze; Stoelting).

**Morris water maze test**

The Morris water maze test was performed as described previously ^30^. Briefly, the acquisition training paradigm for the Morris water maze consisted of eight trials (60 sec maximum; interval 30 min) each day for five consecutive days. Escape latencies, path length, and velocity were recorded during training days. The probe test was performed 24 h after the last acquisition trial. The platform was removed, and mice were introduced into the water from a novel entry point. Mice were allowed to swim freely for 1 minute while the number of platform crossing, time spent in the target quadrant, and latency to the target quadrant were recorded.

**Rotarod test**

The rotarod treadmill (ENV-575, Med Associates, USA) was used to accurately measure motor coordination and fatigue of the cKO and control mice. It consists of a computer-controlled stepper motor-driven drum with constant speed or accelerating speed modes of operation. The apparatus is divided into five test zones so that up to five animals could be tested at the same time. For accelerating speed test, the starting speed was 4 rpm and the speed accelerated from 4.0 to 40 rpm during 300 s. When the animal fell off the rotating drum, it broke a photobeam, leading to the automatic recording of the amount of time spent on the drum and the final speed.

**Open field test**

Open field arenas (MED-VOF-MS, Med Associates, USA) were used to assess anxiety phenotypes and general locomotor activity of the cKO and control mice. This system monitors locomotor activity using video cameras. Videos from the cameras were displayed on the computer screen for user monitoring and recording for later review. Ambulatory and stereotypic behaviors, including ambulatory distance, time spent ambulatory, average velocity, stereotypic time were analyzed by the image analyzing software (ANY-maze; Stoelting).

**Magnetic resonance imaging (MRI)**

Mice were anesthetized with 1% isoflurane. MRI experiments were performed on a 9.4 T/400 mm scanner (Agilent Technologies, Santa Clara, CA), using a quadrature conformal surface coil (7.5mm–12.5mm). Body temperature and respiratory rate were monitored and maintained throughout the experiment. A series of T2-weighted MR images were collected. The T2-weighted images were acquired, typically along coronal orientation, using a RARE (Rapid Acquisition with Relaxation) of Field of view (FOV): 16 mm × 16 mm, matrix size = 192 × 192, slice thickness = 0.5 mm (28 slices, no gap), and bandwidth (BW) = 50 kHz. Acquisition parameters were as follows: repetition time (TR) = 5000 ms, echo spacing = 6.881 ms, Rare factor = 8, effective echo time (TE) = 27.52 ms, flip angle (FA) = 30°, number of averages (NA) = 5. A series of T1-weighted images were acquired using a FLASH (Fast Low Angle Shot) sequence with parameters of FOV: 16 mm × 16 mm, matrix size = 192 × 192, slice thickness = 0.5 mm (28 slices, no gap), BW = 60 kHz, TR = 350 ms, TE = 3 ms, FA = 30°, NA = 5. Brain volumes were analyzed using NIH ImageJ and Matlab programs (MathWorks Inc., Natick, MA).

**Micro-PET scan:**

The radiolabelling synthesis of ^18^F-FDG was conducted as described previously ^31^. Before ^18^F-FDG injection, mice were fasted overnight with free access to water. Mice were injected with approximately 18.5 MBq (500 μCi) of ^18^F-FDG through the tail vein. Thirty minutes later, mice were anesthetized with isoflurane and PET/CT scanning was performed (Siemens Medical Solutions, Malvern, PA), which provided a 12.7 cm axial field of view and an intrinsic resolution < 1.7 ram. The entire static process included CT scanning for 10 min and PET scanning for 20 min. PET images were reconstructed using Fourier rebinning and 2-dimensional filtered back-projection (2D FBP) method (ramp filter and cutoff at Nyquist frequency) with an image matrix of 128 × 128 × 159, resulting in 3D images with a pixel size of 0.77 mm and a slice thickness of 0.796 mm. The images were analyzed using Inveon Research Workplace software (IRW, Siemens Medical Solutions, Malvern, PA). ROIs were automatically extracted from all micro-PET images using ^18^F-FDG murine brain templates of IRW. Relative FDG uptake was calculated for each ROI using the average tissue activity in the region.

**Untargeted Metabolomics**

All of the mice were anesthetized with [pentobarbital](javascript:;) [sodium](javascript:;) and perfused with 4°C PBS to remove blood. The brain samples were quickly taken out from cranial cavity and immediately frozen in liquid nitrogen after the mice were sacrificed. Finally, the brain samples were stored at -80 °C for further use. The measurement would be completed within one month. To cortical samples of the cKO or control mice (100 mg) were added 1 mL cold methanol/acetonitrile/H_2_O（2:2:1, v/v/v）and the mixture was adequately vortexed. The lysate was homogenized by MP homogenizer (24 × 2, 6.0 M/S, 60 s, twice). The homogenate was sonicated on ice (30 min/once, twice) and then centrifuged for 20 min (14000 x g, 4°C). The supernatant was collected and dried in a vacuum centrifuge. For LC-MS analysis, the samples were re-dissolved in 100 μL acetonitrile/water (1:1, v/v). Analyses were performed using a UHPLC (1290 Infinity LC, Agilent Technologies) coupled to a quadrupole time-of-flight (ABSciex Triple TOF 6600). For HILIC separation, samples were analyzed using a 2.1 mm × 100 mm ACQUIY UPLC BEH 1.7 µm column (waters, Ireland). In both ESI positive and negative modes, the mobile phase contained A = 25 mM ammonium acetate and 25 mM ammonium hydroxide in water and B = acetonitrile. The gradient was 85% B for 1 min and was linearly reduced to 65% in 11 min, and then it was reduced to 40% in 0.1 min and kept for 4 min, after which it was increased to 85% in 0.1 min, with a 5 min re-equilibration period employed. For RPLC separation, a 2.1 mm × 100 mm ACQUIY UPLC HSS T3 1.8 µm column (waters, Ireland) was used. In ESI positive mode, the mobile phase contained A = water with 0.1% formic acid and B = acetonitrile with 0.1% formic acid, while in ESI negative mode, the mobile phase contained A = 0.5 mM ammonium fluoride in water and B = acetonitrile. The gradient was 1% B for 1.5 min and was linearly increased to 99% in 11.5 min and kept for 3.5 min. Then it was reduced to 1% in 0.1 min and a 3.4 min of re-equilibration period was employed. The gradients were at a flow rate of 0.3 mL/min, and the column temperatures were kept constant at 25 ℃. A 2 µL aliquot of each sample was injected. The ESI source conditions were set as follows: Ion Source Gas1 (Gas1) as 60, Ion Source Gas2 (Gas2) as 60, curtain gas (CUR) as 30, source temperature: 600 ℃, IonSpray Voltage Floating (ISVF) ± 5500 V. In MS only acquisition, the instrument was set to acquire over the m/z range 60-1000 Da, and the accumulation time for TOF MS scan was set at 0.20 s/spectra. In auto MS/MS acquisition, the instrument was set to acquire over the m/z range 25-1000 Da, and the accumulation time for product ion scan was set at 0.05 s/spectra. The product ion scan was acquired using information dependent acquisition (IDA) with high sensitivity mode selected. The parameters were set as follows: the collision energy (CE) was fixed at 35 V with ± 15 eV; declustering potential (DP), 60 V (+) and −60 V (−); excluding isotopes within 4 Da, candidate ions to monitor per cycle: 10.

The raw MS data (wiff.scan files) were converted to MzXML files using ProteoWizard MSConvert before importing into freely available XCMS software. For peak picking, the following parameters were used: centWave m/z = 25 ppm, peakwidth = c (10, 60), prefilter = c (10, 100). For peak grouping, bw = 5, mzwid = 0.025, minfrac = 0.5 were used. In the extracted ion features, only the variables having more than 50% of the nonzero measurement values in at least one group were kept. Compound identification of metabolites by MS/MS spectra with an in-house database established with available authentic standards. After normalization to total peak intensity, the processed data were uploaded before importing into SIMCA-P (version 14.1, Umetrics, Umea, Sweden), where they were subjected to multivariate data analysis, including Pareto-scaled principal component analysis (PCA) and orthogonal partial least-squares discriminant analysis (OPLS-DA). The 7-fold cross-validation and response permutation testing was used to evaluate the robustness of the model. Metabolites of glycolysis and oxidative phosphorylation with p values less than 0.05 were considered as statistically significant.

**Measurement of TDP, thiamine monophosphate, and thiamine**

Fresh whole blood samples of mice were collected, anticoagulated with heparin, and deproteinized with 7.2% perchloric acid. The brain tissues were homogenized with 100 mM of K_2_HPO_4_ (pH = 5.0) and deproteinized with isometric 6.4% perchloric acid. All samples were centrifuged and supernatants were collected. The levels of TDP, thiamine monophosphate, and thiamine were measured as previously described(*24*).

**Statistical analysis**

Graphpad Prism 7 (version 7.01; GraphPad software) was used for statistical analyses. Student’s t-test for single comparisons or one-way ANOVA for multiple comparisons with appropriate Tukey’s or Dunnett’s Multiple Comparison tests or two-way ANOVA (for comparing two independent variables) followed by Bonferroni’s multiple comparisons test were used to determine statistical differences. Summary results are shown as mean ± SEM.
